# CYK-4 functions independently of its centralspindlin partner ZEN-4 to cellularize oocytes in germline syncytia

**DOI:** 10.1101/196279

**Authors:** Kian-Yong Lee, Rebecca A. Green, Edgar Gutierrez, J. Sebastian Gomez-Cavazos, Irina Kolotuev, Shaohe Wang, Arshad Desai, Alex Groisman, Karen Oegema

**Affiliations:** Ludwig Institute for Cancer Research, Department of Cellular and Molecular Medicine, University of California San Diego, La Jolla, California 92093, USA; Department of Physics, University of California San Diego, La Jolla, California, United States of America

**Keywords:** germline syncytia, intercellular bridge, centralspindlin, MgcRacGAP, CYK-4, RACGAP1, kinesin-6, ZEN-4, ring canal

## Abstract

Throughout metazoans, germ cells undergo incomplete cytokinesis to form syncytia connected by intercellular bridges. Formation of gametes ultimately requires bridge closure. Here, we investigate the contribution of the conserved bridge component centralspindlin to oocyte production in *C. elegans*. Centralspindlin is composed of the Rho family GTPase-activating protein (GAP) CYK-4/MgcRacGAP and the microtubule motor ZEN-4/kinesin-6, which are both essential for cytokinesis. In contrast, we show that oocyte production by the syncytial germline requires CYK-4 but not ZEN-4. Longitudinal imaging after conditional CYK-4 inactivation revealed a role in oocyte cellularization, rather than in generation of syncytial compartments. CYK-4’s lipid-binding C1 domain and the GTPase-binding interface of its GAP domain were individually important for oocyte cellularization and for targeting CYK-4 to bridges, where it contributes to enrichment of active RhoA. These results identify a C1-GAP module in CYK-4 that recruits it to bridges in the germline and directs their closure to produce oocytes.

**IMPACT STATEMENT:** The CYK-4 subunit of centralspindlin, a broadly conserved component of intercellular bridges across metazoa, is required for the cytokinesis-like closure of intercellular bridges that cellularizes oocytes to separate them from germline syncytia.

**MAJOR SUBJECT AREAS:** Cell Biology, Developmental Biology & Stem Cells

## INTRODUCTION

During gametogenesis in the male and female germlines of metazoan species, dividing cells often undergo incomplete cytokinesis to form clusters of cells connected by stable intercellular bridges (Haglund, Nezis, & Stenmark, 2011). In *Drosophila*, where pioneering work on syncytial architecture has been done, germline intercellular bridges are called ring canals (Robinson & Cooley, 1997). The conservation across diverse metazoan species and essential role in fertility (Greenbaum et al., 2006; Haglund et al., 2011; Lei & Spradling, 2016; Robinson & Cooley, 1997) suggests that syncytial germline architecture, in which multiple nuclei reside in compartments that share a common cytoplasm, is important for the generation of viable gametes. The benefits of communal living for germ cell nuclei is an active area of investigation. However, one role for cellular interconnectivity, demonstrated for the female germlines in *Drosophila* and mouse, is to allow the nuclei in adjacent connected cells to serve a nurse function in which they assist in the generation of the large quantities of mRNA, protein, and organelles that are required to properly provision a developing oocyte (King & Mills, 1962; Lei & Spradling, 2016; Pepling, 2016; Robinson & Cooley, 1997).

The stable intercellular bridges in syncytial structures are generated by incomplete cytokinesis, and contain some of the components found at bridges that are resolved by abscission during normal cytokinesis (Haglund et al., 2011). In dividing somatic cells, the contractile ring constricts down around the central spindle, a structure composed of a set of anti-parallel microtubule bundles that forms between segregating chromosomes and concentrates key molecules that promote contractile ring assembly. As constriction completes, the central spindle is converted into a microtubule-based midbody, and the contractile ring is converted into a midbody ring. During normal cell division, the midbody and midbody ring work together to promote abscission, which cuts the intercellular bridge to form the two daughter cells (Agromayor & Martin-Serrano, 2013; Mierzwa & Gerlich, 2014). During the formation of germline cysts, the contractile ring constricts down around the central spindle, which forms a midbody, but abscission does not occur. Instead the midbody is disassembled leaving a stable open intercellular bridge (Dym & Fawcett, 1971; Weber & Russell, 1987).

Ultrastructurally, all intercellular bridges contain an electron dense layer immediately beneath the plasma membrane (Haglund et al., 2011). In the female, but not the male, germline in *Drosophila*, ring canals undergo a maturation process in which they increase in size, recruit additional components and assemble an inner, less electron dense, layer composed of bundled circumferentially organized actin filaments, presumably to facilitate the transfer of cytoplasm and organelles into the developing oocyte (Robinson & Cooley, 1997). As a similar inner layer has not been observed in other systems, it is not yet clear if other types of intercellular bridges employ a similar mechanism for structural reinforcement. When component loading is complete, the intercellular bridges connected to the oocyte presumably need to close to cellularize the egg and separate it from the germline syncytium. Yet, how intercellular bridges/ring canals might be reactivated to constrict in the absence of an intervening central spindle is an interesting question about which little is known.

At a molecular level, two contractile ring components anillin and the septins are often observed, at least transiently in intercellular bridges (Haglund et al., 2011). However, the most strikingly conserved component, observed in all intercellular bridges examined to date, is the centralspindlin complex (Carmena et al., 1998; Greenbaum, Iwamori, Agno, & Matzuk, 2009; Greenbaum, Ma, & Matzuk, 2007; Greenbaum et al., 2006; Haglund et al., 2010; Minestrini, Mathe, & Glover, 2002; Zhou, Rolls, & Hanna-Rose, 2013). Centralspindlin is a heterotetrameric complex composed of a dimer of kinesin-6 (MKLP1 in humans, Pavarotti in *Drosophila*, ZEN-4 in *C. elegans*), and a dimer of a Rho family GTPase-activating protein (CYK4/MgcRacGAP/RACGAP1 in humans, CYK-4 in *C. elegans*, and RacGAP50C/Tum in *Drosophila*) (White & Glotzer, 2012). In *C. elegans*, the CYK-4 N-terminus has a coiled-coil that mediates its dimerization, and a short region prior to the coiled-coil that mediates binding to ZEN-4 (**Figure 1A**; (Mishima, Kaitna, & Glotzer, 2002; Pavicic-Kaltenbrunner, Mishima, & Glotzer, 2007). The CYK-4 C-terminus contains the GAP domain and an adjacent C1 domain that is expected, based on work on human Cyk4 (Lekomtsev et al., 2012), to bind polyanionic phosphoinositide lipids and mediate association with the plasma membrane. ZEN-4 has an N-terminal motor domain and a C-terminal coiled-coil-containing region that mediates its dimerization and association with CYK-4 (**Figure 1A**; (Mishima et al., 2002; Pavicic-Kaltenbrunner et al., 2007). Early in cytokinesis, centralspindlin localizes to and is required for the formation of the central spindle; from this location, it is also thought to play a role in the local activation of the small GTPase RhoA to promote contractile ring assembly (Green, Paluch, & Oegema, 2012; White & Glotzer, 2012). Consistent with a role in activating RhoA, the N-terminus of human Cyk4 is phosphorylated by Plk1 to generate a binding site for the RhoA GEF Ect2 (Burkard et al., 2009; Wolfe, Takaki, Petronczki, & Glotzer, 2009). Late in cytokinesis, the central spindle matures to form the microtubule-based midbody. During this transition, centralspindlin is released from microtubules and concentrates in the midbody ring (Elia, Sougrat, Spurlin, Hurley, & Lippincott-Schwartz, 2011; Green et al., 2013; Hu, Coughlin, & Mitchison, 2012). Important topics of current research include determining how the transition in localization from central spindle microtubules to midbody ring is achieved, and assessing the functional contributions of centralspindlin to the midbody ring and to intercellular bridges in syncytia.

**Figure 1:**
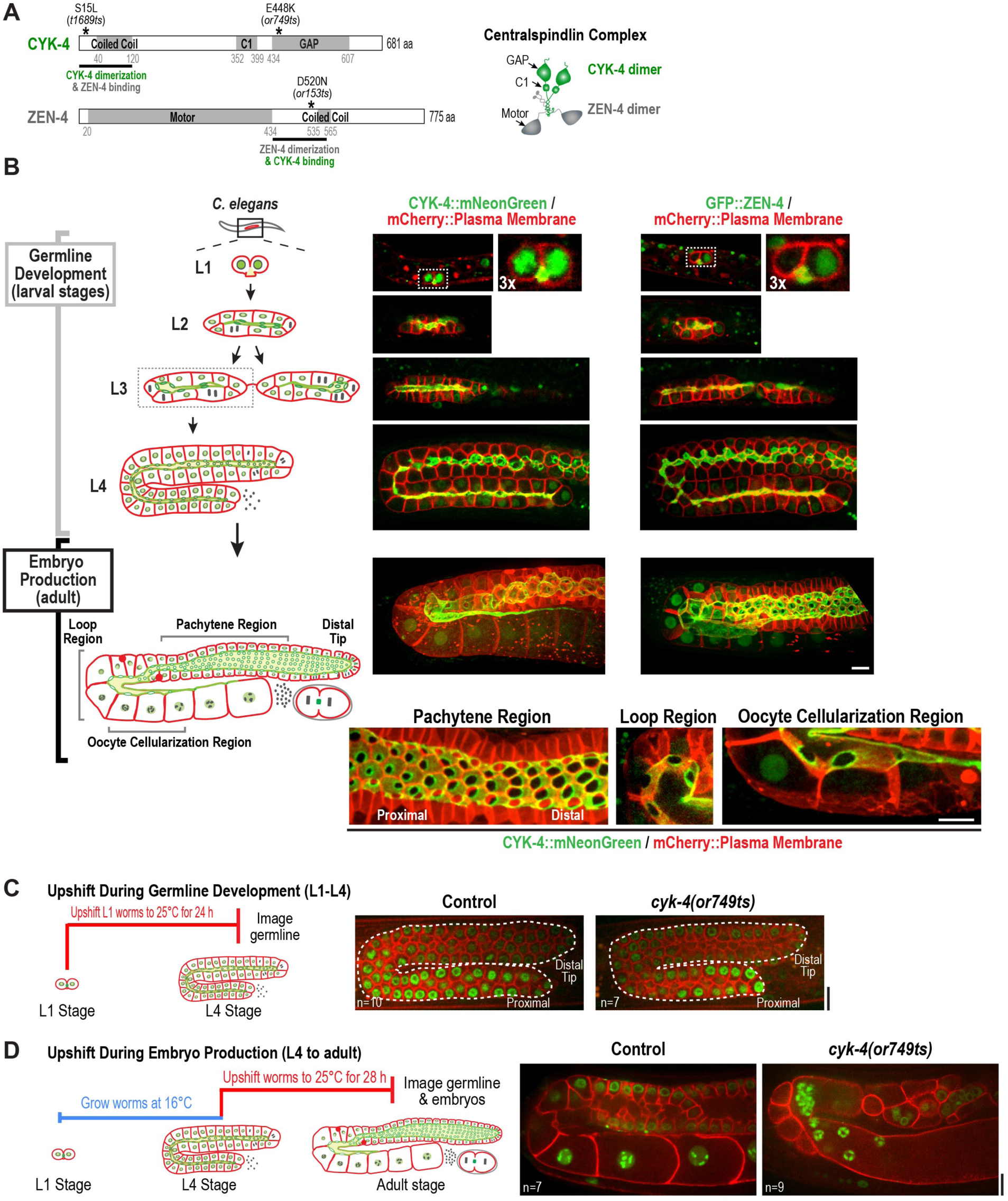
CYK-4 is required for embryo production by the adult germline. (**A**) (*Left*) Schematics highlight the domain structure of the two molecular components, CYK-4 and ZEN-4, of the heterotetrameric centralspindlin complex, and the location of the temperature sensitive mutants used in this study. (*Right*) Schematic showing that centralspindlin is formed by the association of a dimer of CYK-4 with a dimer of ZEN-4. (**B**) (*Left*) Schematics illustrate the development of the syncytial germline. See text for details. (*Right*) Fluorescence confocal images of germlines in worms at the indicated stages expressing an mCherry-tagged plasma membrane marker and either CYK-4::mNeonGreen or *in situ* tagged GFP::ZEN-4. Images for L4 and adult stage germlines are maximum intensity projections. Panels along the bottom row show a higher magnification views of the pachytene, loop, and oocyte cellularization regions in germlines expressing CYK-4::mNeonGreen. Scale bars are 10 μm. (**C**) (*Left*) Schematic outline of the upshift protocol. (*Right*) Representative single plane confocal images of germlines in L4 stage worms (control or *cyk-4(or749ts)*) expressing a GFP-tagged plasma membrane probe (shown in red) and mCherry::histone H2B (shown in green) after the upshift protocol. Dashed lines mark the germline boundaries. (**D**) Schematic outline of the upshift protocol. Representative single plane confocal images of germlines in adult stage worms after the upshift protocol. n in C and D = number of imaged germlines. Scale bars in all panels are 10 μm.

Here, we analyze the role of centralspindlin at intercellular bridges in the *C. elegans* syncytial oogenic germline. During normal cytokinesis, both centralspindlin subunits have an equivalent essential role in central spindle assembly and contractile ring constriction (Canman et al., 2008; Davies et al., 2014; Jantsch-Plunger et al., 2000; Lewellyn, Carvalho, Desai, Maddox, & Oegema, 2011; Loria, Longhini, & Glotzer, 2012; Mishima et al., 2002; Pavicic-Kaltenbrunner et al., 2007; Powers, Bossinger, Rose, Strome, & Saxton, 1998; Raich, Moran, Rothman, & Hardin, 1998; Severson, Hamill, Carter, Schumacher, & Bowerman, 2000). In contrast, we show that in the oogenic germline, CYK-4 is required for the cytokinesis-like event that cellularizes oocytes, severing their connection to the germline syncytium, whereas the kinesin-6 ZEN-4 and its interaction with CYK-4 are not. A domain analysis revealed that the CYK-4 C1 domain and the Rho binding interface of its GAP domain are independently required to recruit CYK-4 to germline bridges and for oocyte cellularization. Consistent with the idea that recruitment to bridges is mediated by binding of the CYK-4 GAP domain to RhoA, knockdown of Rac and Cdc42 did not affect CYK-4 recruitment or oocyte cellularization, whereas RhoA inhibition disrupted germline structure in a fashion similar to CYK-4 inhibition. Cumulatively, our results highlight a critical role for CYK-4 in directing bridge closure to cellularize oocytes, and suggest the C-terminal C1 and GAP domains of CYK-4 form a module required for the targeting of CYK-4 to cortical structures.

## RESULTS

### *Both centralspindlin subunits localize to intercellular bridges throughout* C. elegans *germline development*

The germline in the adult *C. elegans* hermaphrodite (**Figure 1B**, Embryo Production (adult)) consists of ∼1000 nuclei in cup-shaped cellular compartments connected to an extended cytoplasm-containing channel called the rachis (Hall et al., 1999; Hirsh, Oppenheim, & Klass, 1976; Lints & Hall, 2009). The adult germline is a unidirectional pipeline for embryo production, with an oocyte cellularizing to separate from the proximal end of each germline syncytium approximately every 20 minutes. Within the syncytium, mitotic divisions at the distal tip generate new nuclei-containing compartments that progress through the stages of meiotic prophase as they traverse through the pachytene, loop, and oocyte cellularization regions. Transcription in pachytene nuclei produces the material loaded by cytoskeleton-driven flow through the rachis into the oocytes (Gibert, Starck, & Beguet, 1984; Starck & Brun, 1977; Wolke, Jezuit, & Priess, 2007). In the oocyte cellularization region, the compartment bridges close to bud individual oocytes off the syncytium. Oocytes are then fertilized by sperm (produced at an earlier developmental stage) as they pass through the spermatheca to initiate embryogenesis. Consistent with prior work (Zhou et al., 2013), imaging of a functional CYK-4::mNeonGreen fusion (**Figure 1—figure supplement 1**) and an *in situ* tagged GFP::ZEN-4 fusion (**Figure 1— figure supplement 2**) revealed that both proteins localize to the rachis surface (**Figure 1B**), where they are most concentrated around the circular openings into the nuclear compartments (which we will refer to as compartment bridges); lower levels are also detected in nuclei. Compartment bridge diameter decreased by ∼40% during progression through pachytene (from ∼2.5 to 1.4 μm) and then steadily increased concurrent with oocyte loading in the loop region (**Figure 1—figure supplement 3)**. In the oocyte cellularization region, compartment bridge diameter decreased from ∼4 μm to completely closed coincident with tapering of the rachis to a fine tip (**Figure 1—figure supplement 3**).

Monitoring of the two centralspindlin components during development of the germline syncytia (**Figure 1B**, Germline Development (larval stages)) suggested that the rachis is, in fact, an expanded intercellular bridge. The germline arises from a pair of connected primordial germ cells (Z_2_ and Z_3_) that arise from an incomplete cytokinesis in the embryo (Goupil, Amini, & Labbe, 2017; Hubbard & Greenstein, 2005) and remain quiescent until the mid-L1 larval stage. At the L1 larval stage, the intercellular bridge connecting Z_2_ and Z_3_ extends out on one side (**Figure 1B**, **Figure 1—figure supplement 4)**. Examination of serial sections of an L1 germline using electron microscopy revealed that the two nuclei-containing compartments sit side-by-side and open into the small cytoplasm containing intercellular bridge (the nascent rachis; note that the two nuclei-containing compartments exhibit a rounded shape, whereas the intercellular bridge/nascent rachis exhibits a lobed structure outside of the central section (**Figure 1—figure supplement 4; Video 1)**. Subsequent nuclear divisions at the L1 and L2 stages generate additional compartments each with a circular opening to the rachis. At the L3 stage, the germline narrows in the middle, partitioning the syncytium into two arms that increase in length and fold back on themselves during the L4 stage. A dramatic structural reorganization of the germline accompanies the transition to oocyte production in the adult; the rachis increases in width to support component delivery during expansion of the nascent oocytes (**Figure 1B**). Thus, there is a structural continuity through development between the stable intercellular bridge arising from the first incomplete cytokinesis and the rachis in the adult. Consistent with the idea that the rachis is an extended intercellular bridge, the two centralspindlin subunits, which are conserved components of intercellular bridges throughout metazoans, localize to the rachis surface throughout development of the germline syncytia (**Figure 1B**).

### CYK-4 is required for embryo production by the adult germline

To examine centralspindlin function in the germline, we began by using a fast-acting temperature sensitive mutant in the CYK-4 C-terminus (*or749ts*; **Figure 1A**) that is thought to compromise both the GAP and C1 domains at the non-permissive temperature (Canman et al., 2008; Davies et al., 2014; Zhang & Glotzer, 2015). During embryonic cytokinesis, upshift of *cyk-4(or749ts)* embryos leads to a constriction defect similar to CYK-4 depletion (Canman et al., 2008; Davies et al., 2014; Loria et al., 2012; Zhang & Glotzer, 2015). Prior work has also shown that upshift of this C-terminus destabilizing mutant prevents targeting of the mutant protein to the rachis surface in the germline (Zhang & Glotzer, 2015). To separate effects on the proliferation of nuclear-containing compartments during germline development from effects on embryo production in the adult, we used two temperature upshift protocols. To assess the effect on compartment proliferation during germline development, L1 larvae were placed at the non-permissive temperature (25°C) and examined at the L4 stage. Remarkably, despite the penetrant effect of *cyk-4(or749ts)* on cortical remodeling during embryonic cytokinesis, germlines in L1-upshifted animals appeared normal at the L4 stage (**Figure 1C**). Consistent with this, L1 larvae that developed to the L4 stage at 25°C and were then shifted back to the permissive temperature (16°C) laid a normal number of embryos (**Figure 1—figure supplement 5**). In the converse experiment, L4 larvae grown at the permissive temperature were shifted to the non-permissive temperature and analyzed 28 hours later. Under this regime, germline structure was strongly perturbed compared to equivalently treated control worms. All of the partitions in the loop and oocyte cellularization regions were absent and the proximal region of the germline had the appearance of a hollow multinucleated tube (tubulated germline; **Figure 1D**). These results indicate that the CYK-4 C-terminus, which promotes association with the rachis surface, is important for germline morphology and function after the transition to embryo production in the adult but is surprisingly not required to build the germline between the L1 and L4 larval stages.

### ZEN-4 is not essential for embryo production by the adult syncytial germline

The above data show that a mutation in the CYK-4 C-terminus significantly disrupts embryo production by the adult syncytial germline. To determine if this function requires CYK-4 to act in the context of the centralspindlin complex, we depleted CYK-4 or ZEN-4 by injecting dsRNA at the L4 stage and analyzing germline structure after 48 hours. Consistent with the phenotype observed for the *cyk-4(or749ts)* mutant shifted to the non-permissive temperature at the L4 stage (**Figure 1D**), partitions in the loop and oocyte cellularization regions were absent in *cyk-4(RNAi)* worms and the proximal region of the germline had the appearance of a hollow multinucleated tube (**Figure 2A**). As expected for such a severe defect, *cyk-4(RNAi)* worms were unable to produce embryos (**Figure 2B**). However, in striking contrast to CYK-4 depletion, ZEN-4 depletion had very little effect on germline structure (**Figure 2A,C**) or function (**Figure 2B**). We confirmed using *in situ*-tagged GFP::ZEN-4 that the RNAi conditions employed efficiently depleted ZEN-4 from the germline (**Figure 2C**); in addition, *zen-4* RNAi led to 100% embryonic lethality, as expected based on its essential role in embryonic cytokinesis (Mishima et al., 2002; Powers et al., 1998; Raich et al., 1998). These observations suggest that CYK-4 can function in the adult germline independently of ZEN-4. Consistent with this finding, mCherry-tagged CYK-4 (**Figure 2—figure supplement 1**) maintained its localization to the rachis surface and compartment bridges in ZEN-4 depleted worms (**Figure 2D**).

**Figure 2.**
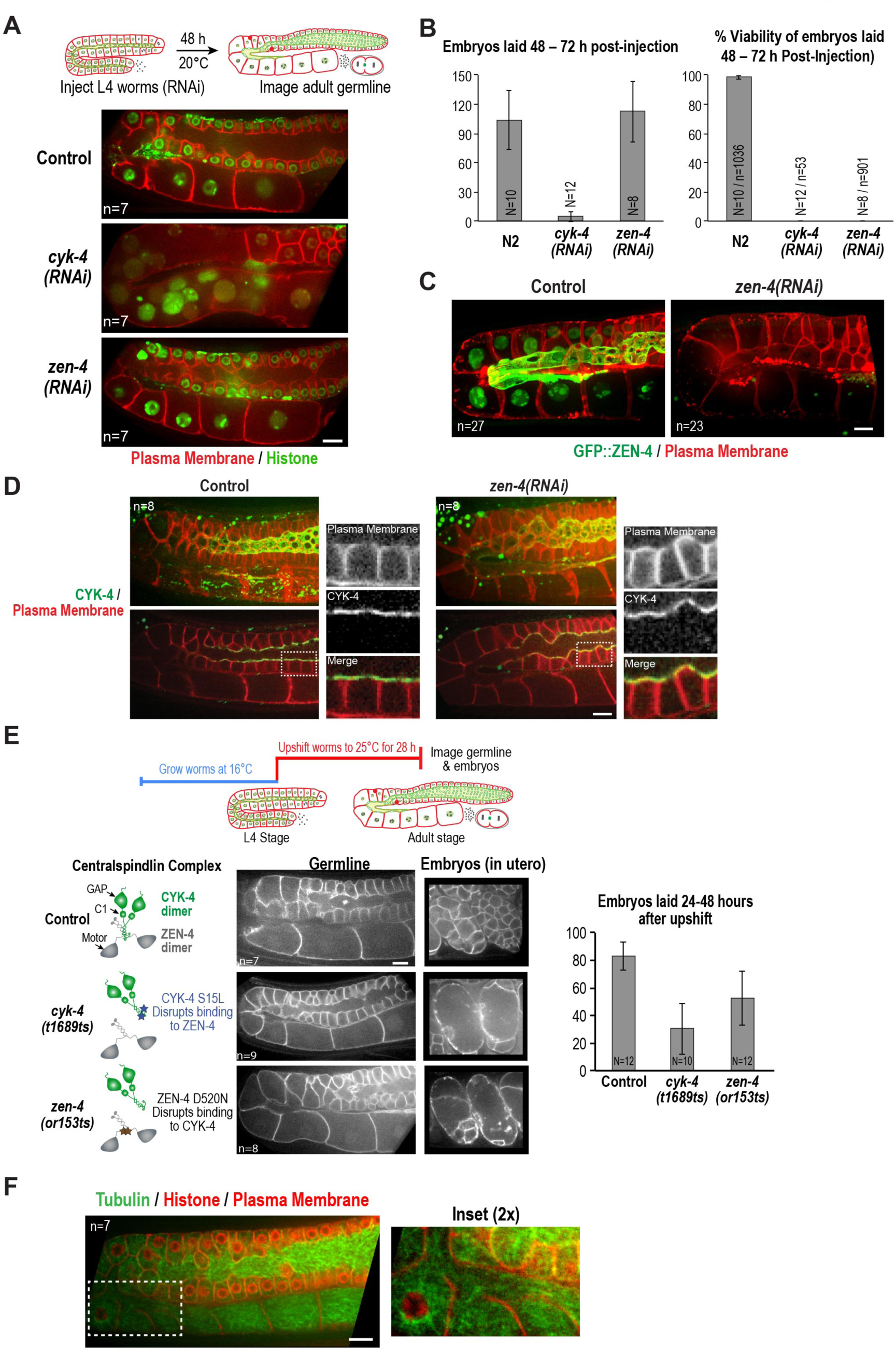
ZEN-4 is not essential for embryo production by the adult syncytial germline. (**A**) (*Top*) Schematic outline of the RNAi experiment. Single plane confocal images of adult germlines expressing a GFP-tagged plasma membrane probe (showninred) and mCherry: histone H2B (shown in green) following depletion of CYK-4 or ZEN-4 by RNAi. n = number of worms imaged. (**B**) Graphs plotting the number of embryos laid during the indicated time intervals (*left*) or embryonic viability (*right*) (mean ± SD) for the indicated conditions. N = number of worms, n = number of embryos. (**C**) Maximum intensity projections of the adult germline in control and *zen-4(RNAi)* worms expressing *in situ* tagged GFP::ZEN-4 and an mCherry-tagged plasma membrane probe. After dsRNA injection, worms were incubated for 48 hours before imaging. n = number of worms. (**D**) Maximum intensity projections (*top*) and single central plane images (*bottom*) of adult germlines in control (*left*) and *zen-4(RNAi)* worms expressing CYK-4::mCherry (shown in green) and a GFP-tagged plasma membrane probe (shown in red). Insets are the boxed region magnified 2.5X. After RNAi injection, worms were incubated for 48 hours before imaging. n = number of imaged worms. (**E**) (*Top*) Schematic outline of the upshift experiment. (*Bottom*, *left*) Single plane confocal images showing the GFP-tagged plasma membrane probe in adult germlines and embryos (in utero) of worms subjected to the upshift protocol. n = number of worms imaged. (*Bottom*, *right*) Graphs plotting the number of embryos laid during the indicated time interval (mean ± SD). N = number of worms. Control data are reproduced from Figure 1—figure supplement 5 for comparison. (**F**) Single plane confocal image of an adult germline in a worm expressing GFP::β-tubulin, mCherry::histone and an mCherry-tagged plasma membrane probe. Inset shows the oocyte cellularization region. n = number of imaged worms. Scale bars in all panels are 10 μm.

To bolster the conclusion that CYK-4 can support germline structure and oocyte production independently of ZEN-4, we examined the effects of two fast-acting temperature sensitive mutants: *cyk-4*(*t1689ts*) and *zen-4*(*or153ts*) that prevent assembly of the heterotetrameric centralspindlin complex (**Figure 1A**; (Encalada et al., 2000; Gonczy et al., 1999; Jantsch-Plunger et al., 2000; Severson et al., 2000)). L4 stage worms were upshifted to the non-permissive temperature (25°C) for 28 hours prior to imaging of the germline and of the embryos in the uterus. Consistent with the RNAi results, germline structure appeared normal and the upshifted mutant worms retained the ability to lay embryos (**Figure 2E**). Notably,penetrant division failure was observed in the embryos present in the *cyk-4*(*t1689ts*) and *zen-4*(*or153ts*) mutants, confirming that the intact centralspindlin complex is essential for cytokinesis (**Figure 2E**).

One reason for why ZEN-4, a microtubule motor, could be required for cytokinesis but not for compartment bridge closure in the germline was suggested by imaging of a strain expressing a GFP fusion with β-tubulin in which the nuclei and plasma membrane were marked with mCherry (**Figure 2F; Figure 2—figure supplement 2**). In contrast to cytokinesis, where contractile rings are bisected by an organized set of anti-parallel microtubule bundles (called the central spindle), we did not observe organized microtubule bundles passing through the compartment bridges in the pachytene, loop, or oocyte cellularization regions of the germline.

We conclude based on both mutant and RNAi analysis that CYK-4 can act independently of ZEN-4 to support embryo production by the syncytial oogenic germline and that this difference may be due to a difference in the role of microtubule-based signaling in the two contexts.

### CYK-4 is required for the intercellular bridge closure that cellularizes oocytes to separate them from the germline syncytium

Our results indicated that depleting CYK-4, or disrupting the C-terminal region containing its GAP and C1 domains using a mutant, leads to a tubulated proximal germline that lacks compartment boundaries (**Figures 1D,2A**). To understand the origin of this phenotype, we developed the means to longitudinally monitor individual living worms over a 5 hour period using a specially made microfluidic device (“worm trap”; **Figure 3A, Figure 3-figure supplement 1**), in which worms are periodically immobilized for imaging and then released to move and feed between timepoints (spaced 50-75 minutes apart). Bacterial suspension is perfused through the microchannel network throughout the imaging session to enable feeding. This setup is conceptually similar to a recently presented microfluidic platform for high-resolution longitudinal imaging of *C. elegans* (Keil, Kutscher, Shaham, & Siggia, 2017), however, the device we designed is simpler and easier to make, because it does not have an embedded microchannel layer. The loading of worms into the device, which is accomplished by manually transferring them into a drop of liquid on top of the device using a standard platinum worm pick, is also straightforward.

**Figure 3.**
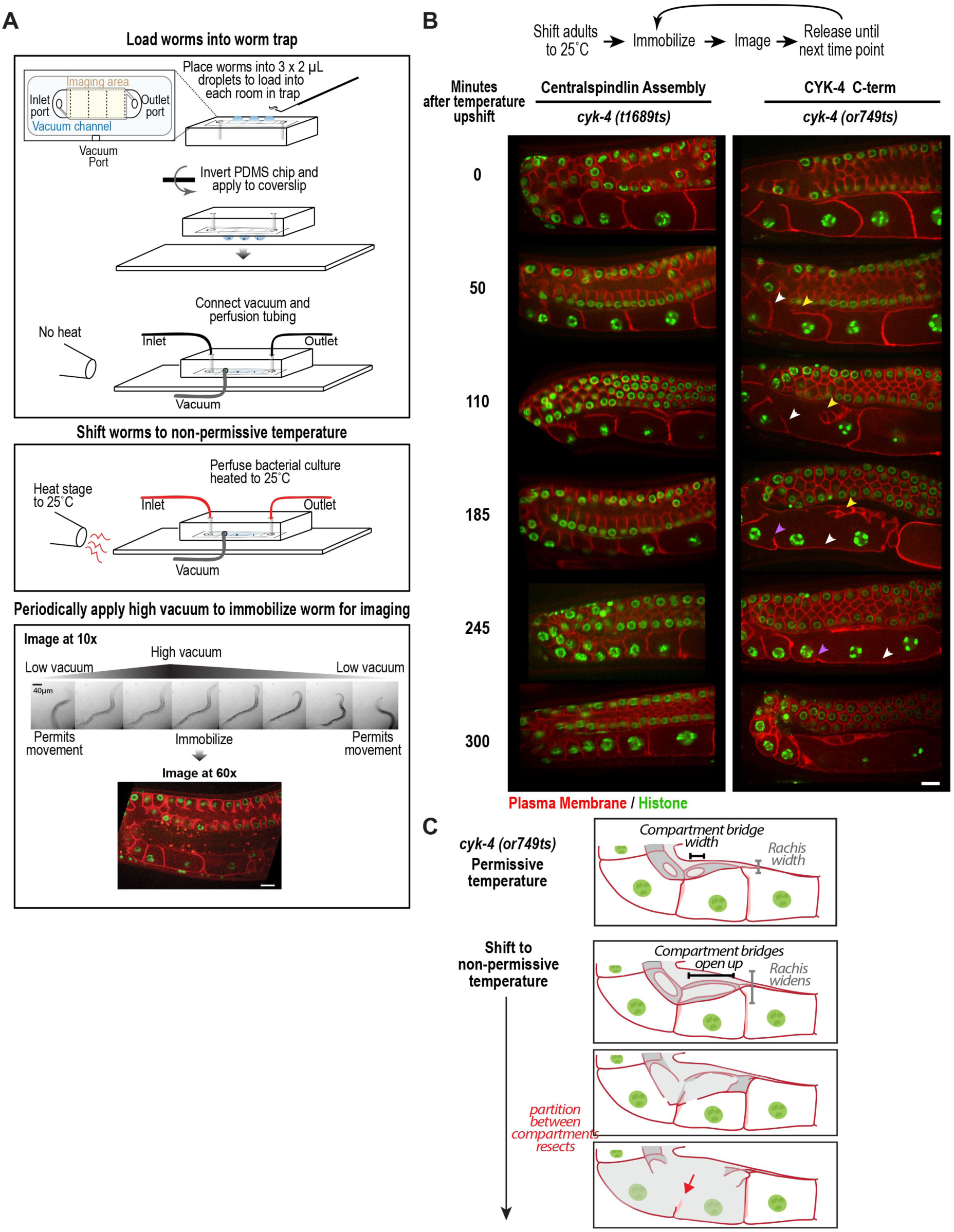
CYK-4 is required for the intercellular bridge closure that cellularizes oocytes to separate them from the germline syncytium. Individual CYK-4 mutant worms were longitudinally monitored using a custom vacuum-actuated microfluidic device (the “worm trap”). (**A**) Schematics summarize the experimental procedure for mounting worms in the trap (*top*), temperature shift (*middle*), and periodic immobilization for imaging (*bottom*). Shown are a series of images demonstrating gradual immobilization and release of a single worm at low magnification (10x), and a single plane high-resolution (60x) image of immobilized worm. (**B**) (*top*) Individual young adult worms carrying the *cyk-4(t1689ts)* mutation that blocks the interaction between CYK-4 and ZEN-4 dimers (n=7; Centralspindlin Assembly) or the *cyk-4(or749ts)* mutation that compromises the CYK-4 C-terminus (n=9; CYK-4 C-term) expressing a GFP-tagged plasma membrane probe (shown in red) and an mCherry fusion with histone H2B (shown in green) were tracked for 5 hours at restrictive temperature in the worm trap. Worms were immobilized periodically for imaging, as indicated in the schematic in (A). Single plane images from the timecourse are shown. (**C**) Schematics illustrate how the phenotype in *cyk-4(or749ts)* worms arises. Following upshift, the partition between the last two uncellularized oocytes resects due to the combined effects of the compartment bridges opening up and the rachis widening (for example, see white arrowheads in the sequence in (B)). Resection of the next and subsequent partitions in a similar fashion (purple arrowheads in the sequence in (B)) leads to a hollow multinucleated “tubulated” proximal germline. Scale bars in all panels are 10 μm.

To determine how the tubulated proximal germline phenotype arises following CYK-4 inhibition, we combined the worm trap imaging approach with temperature upshift of the *cyk-4(or749ts)* mutant that disrupts the C1-GAP domain module in the CYK-4 C-terminus. As a control, we conducted the same analysis with the *cyk-4(t1689ts)* mutant, which disrupts the CYK-4—ZEN-4 interaction but does not significantly affect germline structure or function. Consistent with our single timepoint analysis, upshifting adult *cyk-4(t1689ts)* mutant worms, in which centralspindlin assembly is disrupted, did not prevent oocyte cellularization during the 5-hour imaging period. In contrast, upshifting the *cyk-*4(*or749ts)* mutant, resulted in gradual development of the tubulated germline phenotype (note that this mutant has been previously shown to inactivate CYK-4 within 1 minute during embryonic cytokinesis (Canman et al., 2008)). The phenotype initiated at the proximal tip of the germline where the nascent oocyte was in the process of cellularizing to separate from the germline syncytium. In normal germlines, the diameter of the rachis decreases, tapering to a pointed tip, in parallel with the reduction in the diameter of the compartment bridges that open into the nascent oocytes. The combination of these two events brings the distal partition of the oocyte up to the top of the germline tube and cellularizes the oocyte to bud it off the rachis tip (**Figure 1B**, **Figure 3B**). In *cyk-*4(*or749ts)* germlines after the temperature upshift, the rachis tip above the cellularizing oocyte appeared to widen and lose its structure (**Figure 3B**, yellow arrowheads in 50 and 110 minute timepoints), suggesting that it no longer tapered normally; at the same time, the compartment bridges in the cellularizing oocyte and adjacent compartment opened up (**Figure 3B**, 50 and 110 minute timepoints). Together, these events caused retraction of the partition separating the last uncellularized oocyte from its neighbor (**Figure 3B**, white arrowheads in 50 to 185 minute timepoints, **Figure 3C**). This was followed by opening of the bridge to the next compartment and loss of another partition (purple arrowheads in 185 and 245 minute timepoints).

We conclude, based on longitudinal imaging with a worm trap after temperature upshift, that the germline defect in the *cyk-*4(*or749ts)* mutant arises in a proximal to distal fashion due to failure of the two coordinated events that normally cellularize oocytes: compartment bridge closure and tapering of the rachis. Thus, CYK-4, and specifically its C-terminal C1-GAP module, is critical for the bridge closure that cellularizes oocytes to separate them from the germline syncytium.

### The C1 domain and GAP domain interface that interacts with Rho family GTPases are independently required to target CYK-4 to bridges and for oocyte cellularization

The data above indicate that the C-terminal region of CYK-4 containing the C1 and GAP domains is essential for CYK-4 function in the germline. Prior work has suggested that the CYK-4 C1 domain is required to target CYK-4 to the plasma membrane and that blocking C1 domain-mediated membrane targeting leads to a loss-of-function phenotype (Basant et al., 2015; Zhang & Glotzer, 2015). The *or749ts* mutant potentially disrupts the function of the C1 as well as the GAP domain (Zhang & Glotzer, 2015), precluding assessment of the role of each domain. To analyze the respective roles of the C1 and GAP domains, we therefore generated a set of RNAi-resistant transgenes under the endogenous *cyk-4* promoter encoding untagged wild-type CYK-4 (WT), CYK-4 with the C1 domain deleted (ΔC1), and CYK-4 mutants with changes predicted based on prior structural work to disrupt GAP activity (R459A; mutation of the “arginine finger”) and the ability of the GAP domain to interact with Rho family GTPases (AAE mutant; R459A, K495A and R499E; **Figure 4A**). We note that R459 is part of the GTPase interacting interface, and we cannot exclude the possibility that its mutation could affect GTPase binding as well as GAP activity. All transgenes were expressed at levels comparable to endogenous CYK-4 (**Figure 4B, Figure 4—figure supplement 1**). Examination of germline structure and counting the number of embryos laid 24-48 hours after injection of dsRNA to deplete endogenous CYK-4 confirmed that the transgene encoding WT CYK-4 rescued both germline structure and embryo production. In contrast, the ΔC1 mutant and the R459A and AAE mutants altering residues in the predicted Rho family GTPase interaction interface all failed to rescue (**Figure 4C**). Thus, both the C1 and GAP domains are important for the germline function of CYK-4.

**Figure 4.**
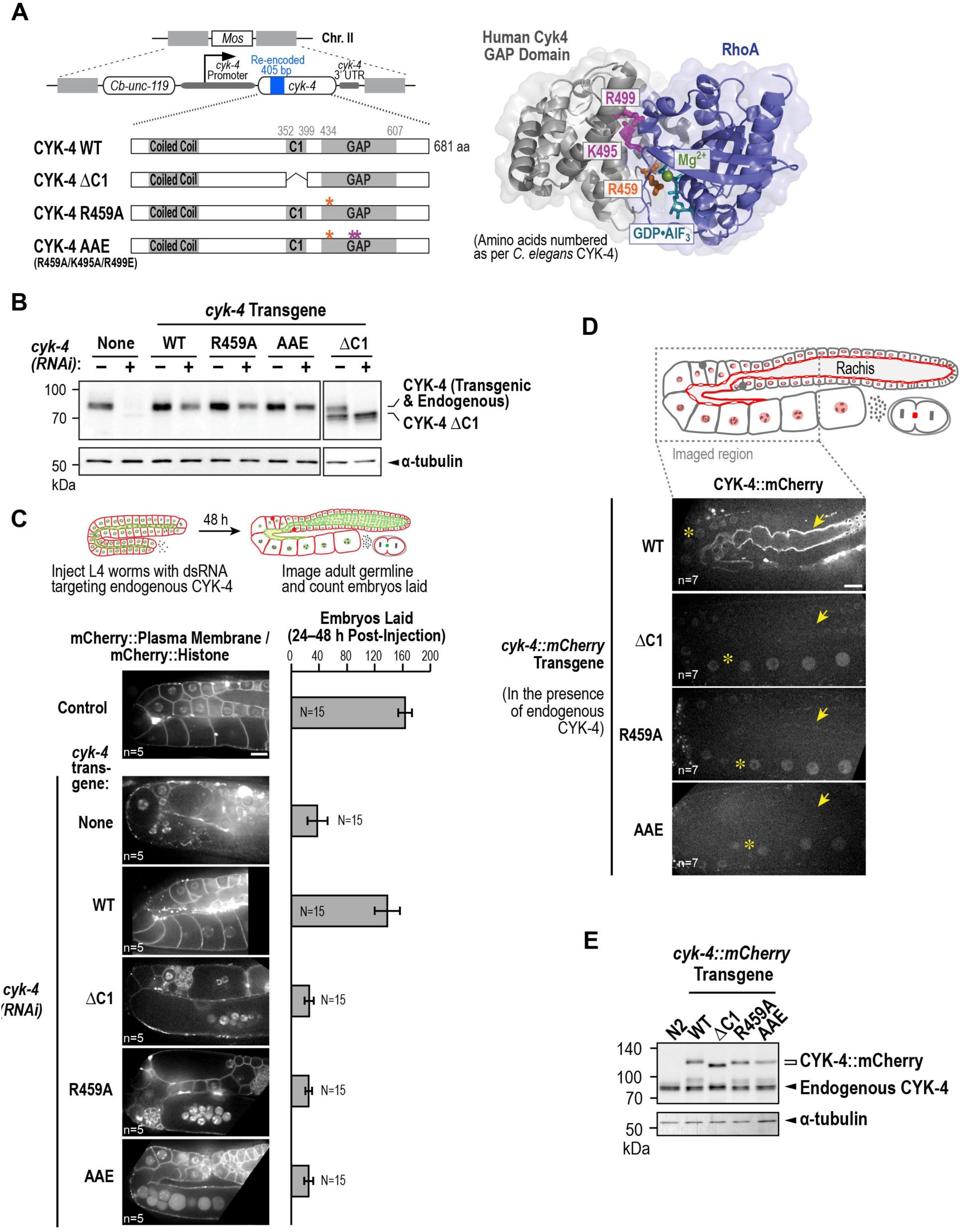
The C1 domain and GAP domain interface that interacts with Rho family GTPases are independently required to target CYK-4 to the rachis surface and for oocyte cellularization. (**A**) (*Left*) Schematic illustrates the set of single copy untagged RNAi-resistant *cyk-4* transgenes inserted into a specific location on Chr. II generated to analyze the role of the C1 and GAP domains. (*Right*) A structural model of the CYK-4 GAP domain (grey; based on the structure of human Cyk4, PDB ID 3W6R) complexed with the Rho family GTPase RhoA (blue; generated by superimposing with ArhGAP20•RhoA complex, PDB ID 3MSX). Key residues in the binding pocket are highlighted in orange (R459, catalytic arginine finger) and magenta (K495 and R499, important for GTPase binding). (**B**) Immunoblots of extracts prepared from worms lacking a transgene (None; N2 strain) or with the transgenes outlined in (A) in the presence (+) or absence (-) of RNAi to deplete endogenous CYK-4. Blots were probed with antibodies to CYK-4 (*top*) and α-tubulin as a loading control (*bottom*). With the exception of the ΔC1 mutant, which runs at a lower molecular weight, the transgene-encoded proteins ran at the same molecular weight as endogenous CYK-4. In the absence of a *cyk-4* transgene, the CYK-4 band disappears, confirming the effectiveness of our RNAi. The protein running at the level of endogenous CYK-4 after RNAi in the WT, R459A, AAE samples is the transgenic protein. (**C**) (*Left*) Single central plane images of the germline in adult worms with the indicated transgenes that were also expressing mCherry::histone and an mCherry plasma membrane maker after depletion of endogenous CYK-4 by RNAi. n = number of worms. (*Right*) Graph plotting the number of embryos laid 24 – 48 hours post-injection (mean ± SD) for strains expressing the indicated *cyk-4* transgenes (without histone or plasma membrane markers). N = number of worms. (**D**) Single central plane images of the germline in adult worms expressing the indicated CYK-4::mCherry fusions. Asterisks highlight the localization of each of the fusions to nuclei and arrows point to the rachis surface in each germline. n = number of imaged worms. (**E**) Immunoblot of extracts prepared from worms expressing the mCherry-tagged RNAi-resistant *cyk-4* transgenes probed with antibodies to CYK-4 (*top*) and α-tubulin (*bottom*) as a loading control. Scale bars are 10 μm.

To examine the effects of the engineered mutations on the ability of CYK-4 to localize to the rachis surface, we generated a parallel series of RNAi-resistant transgenes with a C-terminal mCherry tag. WT CYK-4::mCherry localized to the rachis surface and was enriched in the compartment bridges; it was also present in nuclei, similar to mNeonGreen-tagged CYK-4 (**Figure 4D**, **Figure 1B**). In contrast, the ΔC1, R459A and AAE mutants failed to localize to the rachis surface and compartment bridges, although they were present at levels comparable to the WT protein in nuclei (**Figure 4D**) and localized to central spindles in embryos (*not shown*). Western blotting further confirmed comparable expression of the mCherry-tagged WT and mutant proteins (**Figure 4E**). These results suggest that both the C1 domain and the Rho GTPase binding interface are essential for the targeting of CYK-4 to rachis surface and compartment bridges. We note that these experiments were performed in the presence of endogenous CYK-4, with which the mutant proteins are expected to dimerize via their N-terminal coiled coil, and in the presence of endogenous ZEN-4. Thus, these results suggest that centralspindlin complexes containing even one mutant C1 or one mutant GAP domain are unable to target to the rachis surface. We conclude that the C1 domain that binds lipids and the GAP domain interface that interacts with Rho family GTPases are independently required to target CYK-4 to the rachis surface and for oocyte cellularization.

### Active RhoA localizes to the rachis surface and collaborates with CYK-4 to maintain germline structure

Our results suggested that CYK-4 requires the GAP domain interface predicted to interact with Rho family GTPases to target to the rachis surface and compartment bridges in the germline. We note that in contrast to a prior report (Zhou et al., 2013), depletion of the germline anillin homolog ANI-2, which alters germline structure, did not prevent CYK-4 targeting (**Figure 5—figure supplement 1**). One mechanism by which the GTPase binding interface could contribute to CYK-4 targeting would be interaction with prenylated membrane-anchored GTP-bound RhoA, Rac, or Cdc42. To determine if one of the Rho family GTPase(s) is involved in targeting CYK-4, we measured embryos laid 24-48 hours after injection of dsRNA targeting RhoA (RHO-1 in *C. elegans*), Rac (CED-10 in *C. elegans*; this RNA likely also targets the second *C. elegans* Rac homolog RAC-2) and Cdc42 (CDC-42). Inhibition of Rac^CED-10^ or CDC-42 did not affect embryo production; however, RhoA inhibition led to reduction in embryo production similar to that resulting from CYK-4 depletion (**Figure 5A**). Examination of germline structure following RhoA^RHO-1^ depletion confirmed that it resulted in a tubulated germline phenotype similar to that resulting from depletion of CYK-4 (**Figure 5B**). Imaging germlines expressing mCherry-tagged CYK-4 along with a GFP-tagged plasma membrane probe confirmed that depletion of Rac^CED-10^ or CDC-42 did not affect targeting of CYK-4 to the rachis surface (**Figure 5C**). CYK-4 targeting could not be analyzed following penetrant RhoA^RHO-1^ depletion because the rachis surface is lost in this condition. We note that depletion of Rac^CED-10^ or CDC-42 could not rescue the effects of mutants in the Rho GTPase binding interface on the germline (**Figure 5—figure supplement 2**), suggesting that the critical function of CYK-4 in oocyte cellularization is not to serve as a GAP to reduce levels of active Rac or CDC-42.

**Figure 5.**
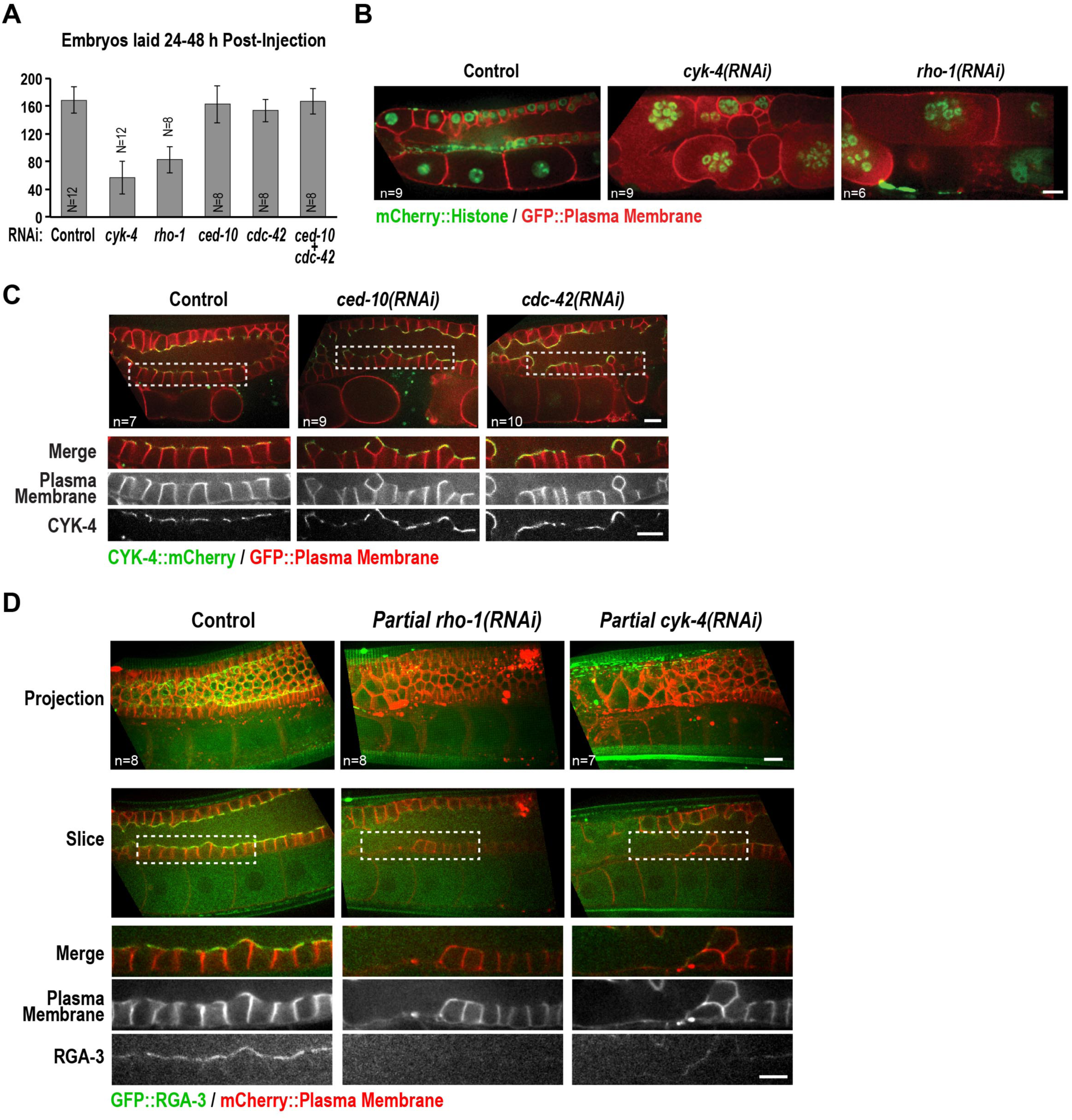
Active RhoA localizes to the rachis surface and collaborates with CYK-4 to maintain germline structure. (**A**) Plot of brood size (mean ± SD) for the indicated RNAi conditions in N2 strain, scored by counting the total number of eggs laid 24 – 48 hours post-injection. N = number of worms. (**B**) Single central plane images of germlines in adult worms expressing mCherry::H2B (shown in green) and a GFP-tagged plasma membrane probe (shown in red) following of CYK-4 or RhoA^RHO-1^ depletion. n = number of worms. (**C**) Single central plane images of germlines in adult worms expressing a GFP-tagged plasma membrane probe (shown in red) and mCherry-tagged CYK-4 (shown in green) following depletion of Rac^CED-10^ or Cdc42^CDC-42^ (48 hours post-injection). Insets (*bottom*) show the boxed regions magnified 1.5x. n = number of worms. (**D**) Maximum intensity projection (*top*) and single central plane images (*middle*) of the germline in adult worms expressing GFP::RGA-3 and an mCherry-tagged plasma membrane probe following partial depletion of RhoA^RHO-1^ or CYK-4 (20 hours post-injection). Insets (*bottom*) show the boxed regions magnified 1.5x. n = number of worms. Scale bars in all panels are 10 μm.

If the Rho GTPase binding interface of the GAP domain targets CYK-4 to the rachis surface by binding to membrane-anchored RhoA, we would expect active RhoA to concentrate in this location. To determine where active RhoA localizes in the germline, we expressed a GFP fusion with RGA-3, a major GAP that targets RhoA in the germline and in embryos (Green et al., 2011; Schmutz, Stevens, & Spang, 2007; Schonegg, Constantinescu, Hoege, & Hyman, 2007; Zanin et al., 2013). Our prior work suggested that RGA-3 localization follows the localization of active RhoA (Zanin et al., 2013). Consistent with this idea, GFP::RGA-3 localized to the rachis surface and this localization was lost following partial inhibition of RhoA under conditions where the structure of the germline was largely intact (**Figure 5D**). This result suggests that active RGA-3-accessible RhoA is concentrated on the rachis surface, supporting the idea that RhoA binding contributes to targeting CYK-4 to this location.

Based on these observations, we propose that both the interaction of the C1 domain with membrane lipids (Lekomtsev et al., 2012) and the interaction of the GAP domain with RhoA are required to recruit CYK-4 to the rachis surface and compartment bridges in the germline (**Figure 6**). Interestingly, partial depletion of CYK-4 under conditions where germline structure remained intact also reduced RGA-3 targeting to the rachis surface (**Figure 5D**). This result suggests existence of a feedback loop in which active RhoA on the rachis surface promotes the recruitment of CYK-4, and CYK-4, in turn, plays a role in generating and/or maintaining active RhoA at this location (**Figure 6**).

**Figure 6.**
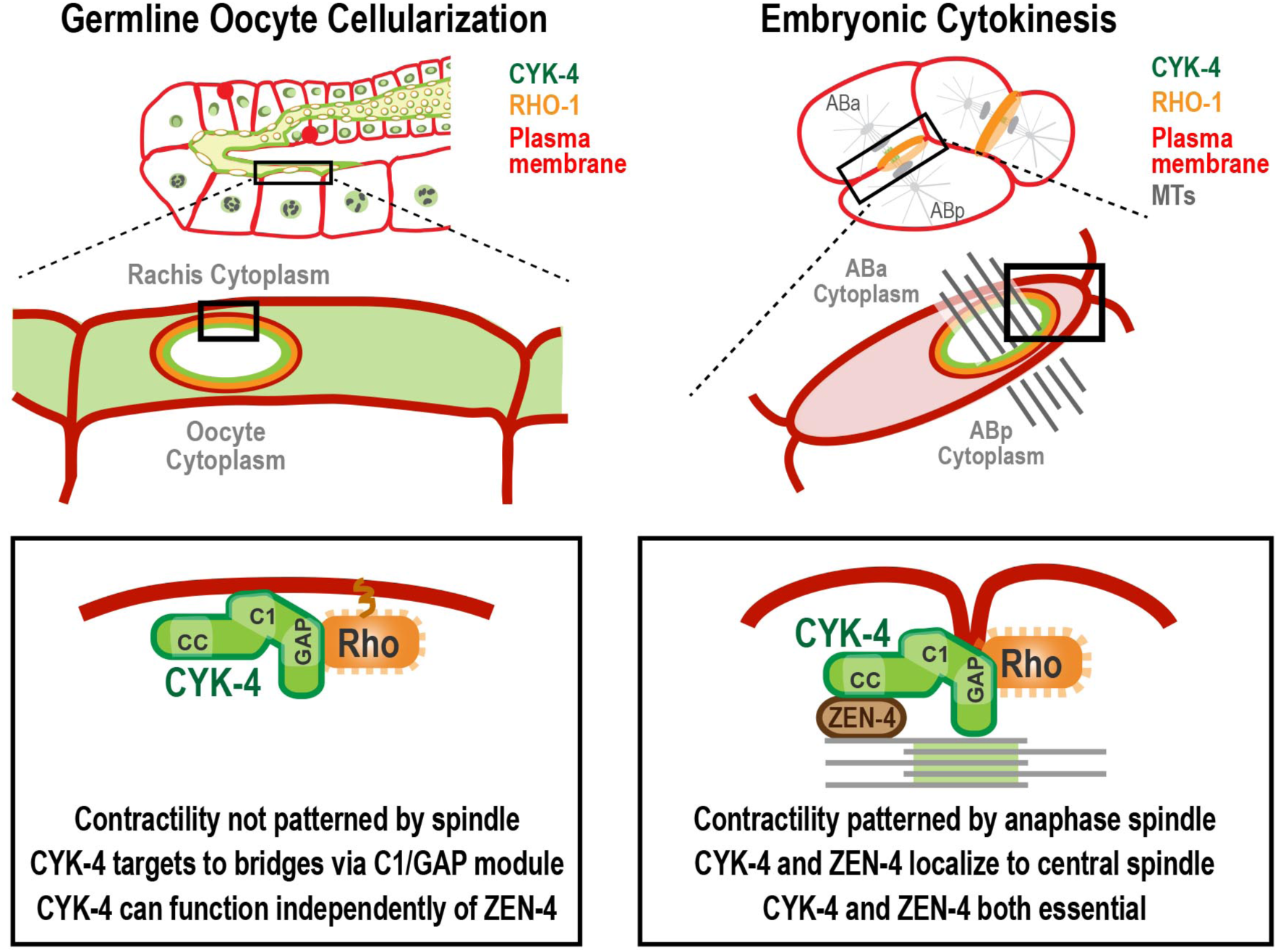
A C-terminal C1 domain-GAP module targets CYK-4 to the rachis surface and compartment bridges to enable oocyte celluarization. Schematics compare the closure of compartment bridges during oocyte cellularization in the germline (*left*) to cytokinesis (shown here in a 2-cell stage embryo, *right*). (*Left*) During bridge closure in the germline, the CYK-4 C1 domain and the Rho GTPase binding interface are both essential for CYK-4 targeting to compartment bridges and the rachis surface. In contrast to cytokinesis, compartment bridges are not bisected by a central spindle and the kinesin-6, ZEN-4, although present, is not essential. (*Right*) During cytokinesis, cortical contractility is patterened by the anaphase spindle. Both ZEN-4 and CYK-4 are required to assemble the central spindle and contractile ring. Targeting to the central spindle and other ZEN-4/microtubule dependent mechanisms may help deliver CYK-4 to the cell surface. However, we speculate that the C-terminal module-based mechanism that operates in the germline may also contribute to CYK-4 delivery during cytokinesis.

## DISCUSSION

During gametogenesis in many metazoans, germ cells undergo incomplete cytokinesis to form syncytia connected by intercellular bridges. In female germlines, this organization facilitates provisioning of the oocyte to support early embryonic development (King & Mills, 1962; Lei & Spradling, 2016; Pepling, 2016; Robinson & Cooley, 1997). Syncytial germ cell development requires a mechanism for closing intercellular bridges to cellularize oocytes prior to fertilization, a process about which very little is known. Here, we use the *C. elegans* oogenic germline as a model to investigate the germline function of centralspindlin, a heterotetrameric complex required for cytokinesis that is also the most conserved component of intercellular bridges in syncytia documented to date (Haglund et al., 2011). Surprisingly, given the equivalent functional importance of the two centralspindlin subunits during cytokinesis, we found that the CYK-4 GAP, but not the ZEN-4 motor, is essential for oocyte production. Using a conditional temperature sensitive mutant, we show that CYK-4 is not required for germline development, but becomes essential during the transition to the adult, when the germline begins to make oocytes. Longitudinal imaging after conditional CYK-4 inactivation using a specially designed microdevice revealed that CYK-4 is specifically required for the cytokinesis-like intercellular bridge closure that cellularizes developing oocytes to separate them from the germline syncytium. Engineered mutants revealed that the lipid-binding C1 domain and the Rho GTPase binding interface of the CYK-4 GAP domain are individually essential to localize CYK-4 to bridges and for oocyte cellularization. Cumulatively, these results provide molecular insight into the poorly understood but widely conserved process of oocyte cellularization, and highlight a critical, kinesin-6-independent role in this process for the Rho GAP subunit of centralspindlin. They also identify a new role for the Rho GTPase-binding interface of the GAP domain in targeting CYK-4 to cortical structures that may be relevant in contexts outside of oocyte cellularization.

### *The* C. elegans *rachis: an expanded intercellular bridge*

In *Drosophila* and mice, germ cells undergo multiple rounds of incomplete cytokinesis, to generate germline cysts in which cells are connected to one or more additional cells by stable intercellular bridges. One (*Drosophila*) or several (mice) of the most connected cells in the cyst differentiate to become oocytes and the other cells serve a nurse-like function, transferring cytoplasm and organelles to the developing oocyte(s) prior to undergoing programmed cell death (Jenkins, Timmons, & McCall, 2013; Lei & Spradling, 2016; Robinson & Cooley, 1997). The oogenic *C. elegans* germline is organized in a more egalitarian fashion; germ cell compartments in the pachytene region contribute material to the expanding compartments in the loop region, and are later loaded with components contributed by the compartments following behind them (Gibert et al., 1984; Hall et al., 1999; Hirsh et al., 1976; Lints & Hall, 2009; Starck & Brun, 1977; Wolke et al., 2007). This assembly line-like progression is accompanied by a variation in syncytial architecture; instead of a network of cells that open through intercellular bridges into each other, all of the cells in the *C. elegans* germline open through bridges into a common cytoplasmic channel called the rachis (Hall et al., 1999; Hirsh, Oppenheim, & Klass, 1976; Lints & Hall, 2009).

In the *C. elegans* oogenic germline, centralspindlin localizes to the rachis surface as well as the compartment bridges (**Figure 1, Figure 1—figure supplements 1-3**; (Zhou et al., 2013)). Our live fluorescence and EM analysis of the nascent germline suggests that this is because the rachis is an expanded intercellular bridge that arises from the first incomplete division of the germline precursor. As the germline increases in size during development, subsequent nuclear divisions generate additional compartments while remaining connected to this expanded bridge. Consistent with the rachis being an extended intercellular bridge, all of the components found on the compartment bridges, including the anillin homolog ANI-2 (Amini et al., 2014; Maddox,Habermann, Desai, & Oegema, 2005) and the BTB domain containing protein CYK-7 ((Green et al., 2011); our unpublished results) are also enriched on the rachis surface. The RhoA GAP RGA-3 also specifically targets to the rachis surface (**Figure 5**), suggesting that active RhoA is concentrated at this location. Other components such as myosin II, ANI-1 and the septins localize to the rachis surface and compartment bridges, but are also found along the sides of the compartments (Maddox et al., 2005). The similar composition of the rachis surface and compartment bridges is consistent with the concurrent constriction of the rachis and the compartment bridges during oocyte cellularization.

### Regulation of oocyte cellularization

The microfluidic device-assisted longitudinal imaging of *cyk-4(or749ts)* mutant worms indicates that CYK-4 controls the simultaneous constriction of the rachis and compartment bridges to cellularize oocytes. An important question raised by these findings is what triggers these events specifically in the oocyte cellularization region of the germline. During cytokinesis in dividing cells, the key regulatory event is reduction of Cdk1 kinase activity due to the activation of the anaphase-promoting complex/cyclosome (APC/C), the E3 ubiquitin ligase that targets cyclin B for degradation (Green et al., 2012). The cytokinesis-like event that cellularizes oocytes occurs near the end of meiotic prophase, prior to nuclear envelope breakdown and the first round of meiotic chromosome segregation. How or whether cell cycle regulators bring about oocyte expansion and cellularization is an open question about which relatively little is known. Consistent with the idea that oocyte cellularization could be triggered by APC/C activation, we identified a phenotypic class in a screen of proteins required for embryo production that contained APC/C subunits and was characterized by the accumulation of oocytes with large open compartment bridges and compromised rachis tapering (Green et al., 2011). More work will be needed to determine whether transient APC/C activation is indeed the trigger for cortical constriction during oocyte cellularization, as it is for contractile ring assembly and constriction during cytokinesis. If the APC/C controls this transition, it will also be interesting to determine whether the key target is a cyclin as it is during mitotic exit. We note that MAPK signaling, which is active in the pachytene and oocyte cellularization regions, but not the intervening loop region, is repressed by cyclin A in the loop region, (Arur et al., 2009). This observation suggests cyclin A may be active in the loop region and inactivated in the oocyte cellularization region, a possibility that should be addressable in future work.

### Potential reasons why CYK-4 can function independently of ZEN-4 in the germline

Our data indicate that CYK-4 can function independently of ZEN-4 in oocyte cellularization. Although CYK-4 localizes to the rachis surface/compartment bridges and supports oocyte cellularization independently of ZEN-4, both proteins are normally present. These results raise the question of why CYK-4 can function independently of ZEN-4 in the germline, which is in striking contrast to cytokinesis. One possibility is that different mechanisms drive association of CYK-4 with the cortex in the germline versus in embryos. Centralspindlin must go to the cortex to function in both contexts based on its proposed roles in activating RhoA and/or inactivating Rac (Green et al., 2012; Loria et al., 2012; White & Glotzer, 2012; Zhang & Glotzer, 2015). In addition, the lipid-binding C1 domain of *C. elegans* CYK-4 is essential in both contexts ((Zhang & Glotzer, 2015); **Figure 4C**). However, there is a significant difference in the amount of centralspindlin associated with the cortex in the germline versus during embryonic cytokinesis. In the germline, centralspindlin localizes prominently to the rachis surface/compartment bridges. In contrast, during cytokinesis its most prominent localization is to microtubules and/or the overlapping microtubule bundles of the central spindle (weak cortical localization has been reported during cytokinesis using CYK-4::GFP expressed with an exogenous *pie-1* 3’ UTR that potentially drives overexpression (Basant et al., 2015; Zhang &Glotzer, 2015); in contrast, using functional transgenes with endogenous *cyk-4* promoter and 3’ UTR we have failed to detect CYK-4 at the cortex during cytokinesis in early embryos (*not shown*)). A second major difference is that during cytokinesis, targeting of centralspindlin to the cortex is proposed to be controlled by its localization to anti-parallel microtubule bundles in the central spindle (White & Glotzer, 2012) and by ZEN-4 motor-based movement along and accumulation at the ends of stabilized astral microtubules (Breznau, Murt, Blasius, Verhey, & Miller, 2017; Foe & von Dassow, 2008; Nishimura & Yonemura, 2006; Odell & Foe, 2008; Vale, Spudich, & Griffis, 2009). In contrast, in the region of the germline where oocyte cellularization occurs there are no centrosomes (Mikeladze-Dvali et al., 2012) and no central spindle-like microtubule bundles (**Figure 2F**). Thus, CYK-4 may function independently of ZEN-4 in the germline because association of CYK-4 with the cortex is negatively regulated during cytokinesis to ensure its control by microtubule-based signaling. Alternatively, association of CYK-4 with the cortex may be positively regulated in the germline by a modification or protein-protein interaction that is absent in dividing embryos. Additional work will be needed to discriminate between these two potential mechanisms.

### The Rho GTPase binding interface of the GAP domain and the C1 domain are independently critical for localizing CYK-4 in the germline

The prominent cortical localization of CYK-4 in the germline allowed us to analyze the elements in CYK-4 required for this localization and correlate with function in oocyte cellularization. The CYK-4 C-terminus has previously been shown to be essential for targeting CYK-4 to the cortex in the germline and in embryos (Basant et al., 2015; Zhang & Glotzer, 2015), although the functional element required for targeting was assumed to be the C1 domain (Lekomtsev et al., 2012). We find that the Rho GTPase-binding interface of the GAP domain and the C1 domain are both independently required for localizing CYK-4 to the rachis surface and for oocyte cellularization. Based on the fact that active RhoA is enriched on the rachis surface, we propose that the Rho GTPase-binding interface of the CYK-4 GAP domain interacts with membrane-associated RhoA, and that this interaction collaborates with binding of the C1 domain to phosphoinositide lipids (**Figure 6**).

We found that CYK-4 mutants with alterations in the C1 domain or the GTPase binding interface were unable to target even in the presence of endogenous CYK-4. These results imply that CYK-4, which dimerizes via its N-terminal coiled-coil, requires two functional C1 domains and two functional Rho GTPase binding interfaces for robust localization. We note that the mechanism that we propose here is reminiscent of the mechanism recently proposed based on structural work for the membrane association of the contractile ring protein anillin, which requires three low affinity membrane-associating elements, a cryptic C2 domain, a RhoA-binding domain and a PH domain (Sun et al., 2015).

In summary, our results provide molecular insight into how oocytes are cellularized to separate from germline syncytia. Cellularization requires the CYK-4 subunit of centralspindlin but does not require its binding partner ZEN-4. Cellularization is critically dependent on its C-terminal C1 and GAP domains of CYK-4, which are both essential for its recruitment to intercellular bridges. These findings raise questions about the regulation of oocyte cellularization and the mechanisms that direct RhoA generation and contractility in the germline and how these relate to centralspindlin function during cytokinesis that will be interesting topics for future work.

## MATERIALS AND METHODS

### *C. elegans* strains

*C. elegans* strains (listed in **Table 1**) were maintained at 20°C, except for strains containing temperature sensitive alleles, which were maintained at 16°C. Transgenes generated for this study were inserted in single copy at specific chromosomal sites using the transposon-based MosSCI method (Frokjaer-Jensen et al., 2008). Depending on which Mos1 insertion site was used, transgenes were cloned into pCFJ151 (*ttTi5605* site on Chr II; Uni I *oxTi185* site on Chr I; Uni IV *oxTi177* site on Chr IV) or pCFJ352 (*ttTi4348* site on Chr I). Transgenes were generated by injecting a mixture of repairing plasmid containing the *Cb-unc-119* selection marker and appropriate homology arms (50 ng/μL), transposase plasmid (pCFJ601, P*eft-3::Mos1 transposase*, 50 ng/μL) and four plasmids encoding fluorescent markers for negative selection [pMA122 (P*hsp-16.41::peel-1*, 10 ng/μL), pCFJ90 (P*myo-2::mCherry*, 2.5 ng/μL), pCFJ104 (P*myo-3::mCherry*, 5 ng/μL) and pGH8 (P*rab-3::mCherry*, 10 ng/μL)] into strains EG6429 (outcrossed from EG4322; *ttTi5605*, Chr II), EG6701 (*ttTi4348*, Chr I), EG8078 (*oxTi185*, Chr I) or EG8081 (*oxTi177*, Chr IV). After one week, progeny of injected worms were heat-shocked at 34°C for 2 hours to induce the expression of PEEL-1 to kill extra chromosomal array containing worms (Seidel et al., 2011). Moving worms without fluorescent markers were identified as candidates, and transgene integration was confirmed in their progeny by PCR across the junctions on both sides of the integration site.

**Table 1.** *C. elegans* strains used in this study

| Strain # | Genotype | Figure |
| --- | --- | --- |
| N2 | wild type (ancestral) | 1S1, 1S4B, 2B, 2S1, 4B, 4C, 4E, 4S1, 5A, 5S2 |
| OD95 | *unc-119(ed3) III; ItIs37 [pAA64; Ppie-1::mCherry::his-58; unc-119 (+)] IV; ItIs38 [pAA1; Ppie-1::GFP::PH(PLC1delta1); unc-119(+)] III* | 1C, 1D, 1S5, 2A, 2E, 5B, 1S4A |
| OD239 | *cyk-4(or749ts) III; ItIs38 [pAA1; Ppie-1::GFP::PH(PLC1delta1) unc-119 (+)] III; ItIs37 [pAA64; Ppie-1::mCherry::H2B his-58; unc-119 (+)] IV* | 1C, 1D, 1S5, 3B, 2E |
| OD241 | *cyk-4(t1689ts) III; ItIs38 [pAA1; Ppie-1::GFP::PH(PLC1delta1) unc-119 (+)] III; ItIs37 [pAA64; Ppie-1::mCherry::H2B his-58; unc-119 (+)] IV* | 2E, 3B |
| OD1176 | *unc-119(ed3) III; ItSi346 [pKL3; Pcyk-4::CYK-4reencoded::mCherry; cb-unc-119(+)] IV* | 2S1, 4D, 4E |
| OD1178 | *unc-119(ed3) III; ItSi348 [pKL4; Pcyk-4::CYK-4reencoded(R459A)::mCherry; cb-unc-119(+)] IV* | 4D, 4E |
| OD1211 | *ItSi346 [pKL3; Pcyk-4::CYK-4reencoded::mCherry; cb-unc-119(+)] IV; ItIs38 [pAA1; Ppie-1::GFP::PH(PLC1delta1); unc-119 (+)] III* | 2D, 5C |
| OD1364 | *unc-119(ed3) III; ItSi432[pKL33; Pcyk-4::CYK-4reencoded(ΔC1)::mCherry; cb-unc-119(+)] IV* | 4D, 4E |
| OD1970 | *ItSi835 [pKL62; Pcyk-4::CYK-4reencoded; cb-unc-119(+)]II; unc-119(ed3) III* | 4B, 4S1, 5S2 |
| OD1972 | *ItSi837 [pKL64; Pcyk-4::CYK-4reencoded(R459A); cb-unc-119(+)] II; unc-119(ed3) III* | 4B, 4S1, 5S2 |
| OD1974 | *ItSi839 [pKL65; Pcyk-4::CYK-4reencoded(R459A/K495A/R499E); cb-unc-119(+)] II; unc-119(ed3) III* | 4B, 4S1, 5S2 |
| OD1978 | *ItSi843 [pKL67; Pcyk-4::CYK-4reencoded(ΔC1); cb-unc-119(+)] II; unc-119(ed3) III* | 4B, 4S1 |
| OD2064 | *ItSi849[pKL120; Pmex-5::mCherry-PH::tbb-2 3'UTR; cb-unc-119(+)]I; ItSi641[pKL89; Pcyk-4::CYK-4reencoded::GFP::MEI-1(1–224); cb-unc-119(+)]I; unc-119(ed3)III; ItIs37[pAA64; pie-1/mCherry::H2B his-58; unc-119(+)] IV* | 4C |
| OD2083 | *ItSi849 [pKL120; Pmex-5::mCherry::PH(PLC1delta1)::tbb-2 3'UTR; cb-unc-119(+)] I; ItSi641 [pKL89; Pcyk-4::CYK-4reencoded::GFP::MEI-1(1–224); cb-unc-119(+)] I; ItSi835 [pKL62; Pcyk-4::CYK-4reencoded; cb-unc-119(+)]II; unc-119(ed3) III; ItIs37 [pAA64; Ppie-1::mCherry::H2B his-58; unc-119(+)] IV* | 4C |
| OD2084 | *ItSi849 [pKL120; Pmex-5::mCherry:: PH(PLC1delta1)::tbb-2 3'UTR; cb-unc-119(+)] I; ItSi641 [pKL89; Pcyk-4::CYK-4reencoded::GFP::MEI-1(1–224); cb-unc-119(+)] I; ItSi837 [pKL64; Pcyk-4::CYK-4reencoded(R459A); cb-unc-119(+)] II; unc-119(ed3) III; ItIs37 [pAA64; Ppie-1::mCherry::H2B his-58; unc-119(+)] IV* | 4C |
| OD2085 | *ItSi849 [pKL120; Pmex-5::mCherry::PH(PLC1delta1)::tbb-2 3'UTR; cb-unc-119(+)] I; ItSi641 [pKL89; Pcyk-4::CYK-4reencoded::GFP::MEI-1(1–224); cb-unc-119(+)] I; ItSi839 [pKL65; Pcyk-4::CYK-4reencoded(R459A/K495A/R499E); cb-unc-119(+)] II; unc-119(ed3) III; ItIs37 [pAA64; Ppie-1::mCherry::H2B his-58; unc-119(+)] IV* | 4C |
| OD2087 | *ItSi849 [pKL120; Pmex-5::mCherry::PH(PLC1delta1)::tbb-2 3'UTR; cb-unc-119(+)]I; ItSi641 [pKL89; Pcyk-4::CYK-4reencoded::GFP::MEI-1(1-224); cb-unc-119(+)] I; ItSi843 [pKL67; Pcyk-4::CYK-4reencoded(ΔC1); cb-unc-119(+)] II; unc-119(ed3) III; ItSi37 [pAA64; pie-1::mCherry::H2B his-58; unc-119(+)] IV* | 4C |
| OD2127 | *ItSi220 [pOD1249/pSW077; Pmex-5::GFP-tbb-2-operon-linker-mCherry-his-11; cb-unc-119(+)] I; ItSi849 [pKL120; Pmex-5::mCh- PH::tbb-2 3'UTR; cb-unc-119(+)] I* | 2F, 2S2 |
| OD2286 | *unc-119(ed3) III; ItSi867 [pKL142; Pcyk-4::CYK-4reencoded(R459A/K495A/R499E)::mCherry; cb-unc-119(+)] IV* | 4D, 4E |
| OD3639 | *ItSi849 [pKL120; Pmex-5::mCherry::PH(PLC1delta1)::tbb-2 3'UTR; cb-unc-119(+)] I; ItSi17 [pOD928/EZ-36; Prga-3::GFP::RGA-3; cb-unc-119(+)] II; unc-119(ed3) III* | 5D |
| OD3640 | *ItSi849 [pKL120; Pmex-5::mCherry::PH(PLC1delta1)::tbb-2 3'UTR; cb-unc-119(+)] I; unc-119(ed3) III(?); zen-4(It30[GFP::loxP::zen-4]) IV* | 1B, 1S2, 2C |
| OD3686 | *ItSi849 [pKL120; Pmex-5::mCherry::PH(PLC1delta1)::tbb-2 3'UTR; cb-unc-119(+)] I; ItSi1124[pSG092; Pcyk-4::CYK-4reencoded::mNeonGreen::cyk-4 3'-UTR; cb-unc-119(+)] II; unc-119(ed3) III* | 1B, 1S1, 1S3, 5S1 |
| JCC754 | *unc-119(ed3) III?; ItSi38 [pAA1; Ppie-1::GFP::PH(PLC1delta1); unc-119 (+)] III; zen-4(or153ts)IV; ItSi37 [pAA64; Ppie-1::mCherry::his-58; unc-119 (+)] IV* | 2E |

**Table 2.**
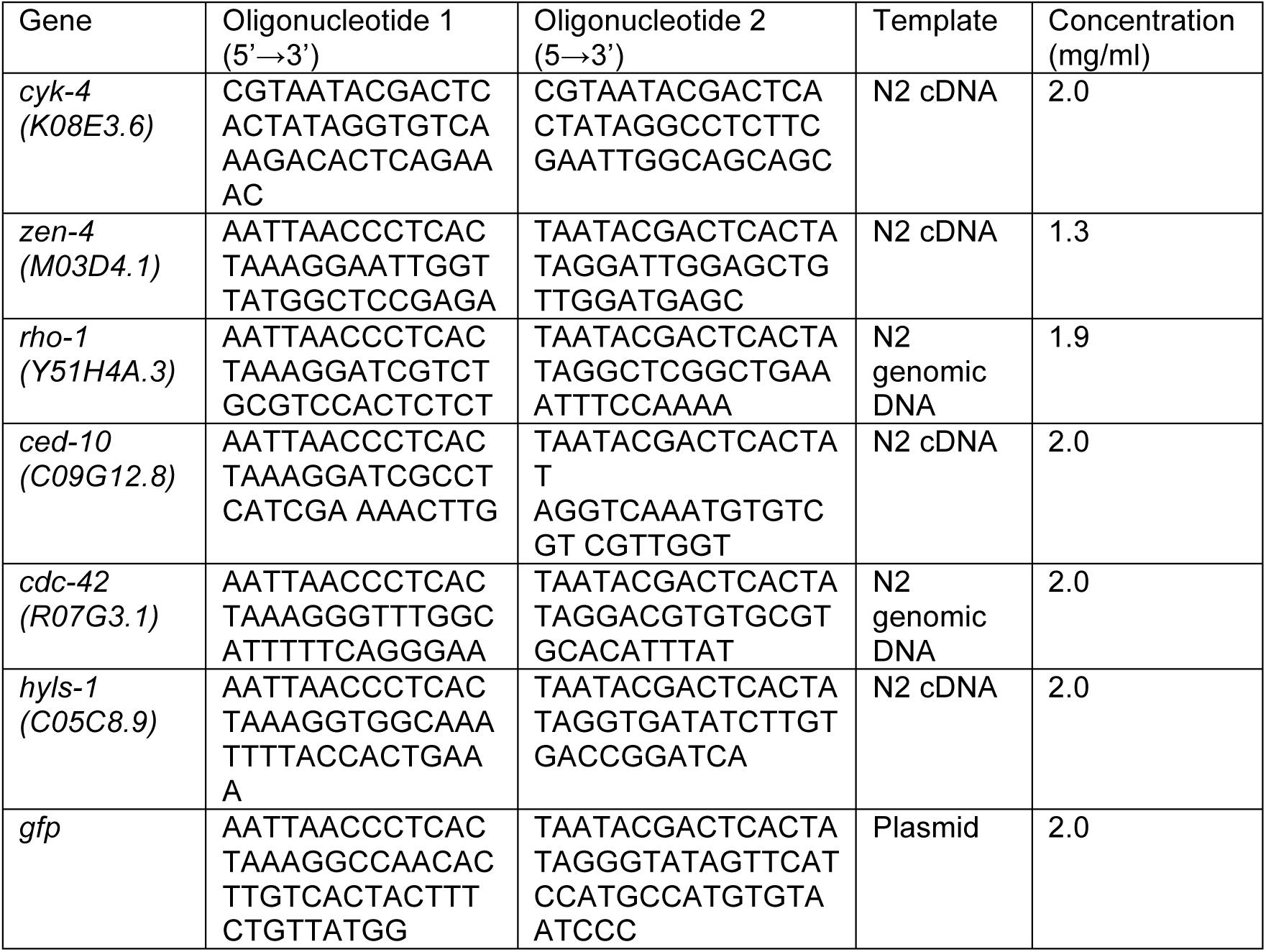
Oligos used for dsRNA production

A CRISPR/Cas9-based method (Dickinson, Pani, Heppert, Higgins, & Goldstein, 2015) was used to generate an *in situ*-tagged *gfp::zen-4* strain. Briefly, a repairing plasmid containing a loxP-flanked self-excision cassette (SEC; containing a dominant roller marker, a heat-shock inducible Cre recombinase and a hygromycin drug resistance marker) and GFP, together surrounded by homology arms for N-terminal insertion at the *zen-4* locus (651 bp of the *zen-4* 5’UTR and 541 bp of the *zen-4* coding sequence; pOD2083, 10 ng/μL) was co-injected with a plasmid encoding the Cas9 protein modified from pDD162 by inserting a guide RNA sequence (5’-AAATGTCGTCGCGTAAACG-3’, 50 ng/μL) and three plasmids encoding fluorescent markers for negative selection [pCFJ90 (P*myo-2::mCherry*, 2.5 ng/μL), pCFJ104 (P*myo-3::mCherry*, 5 ng/μL) and pGH8 (P*rab-3::mCherry*, 10 ng/μL)] into the wildtype N2 strain. After a week, roller worms without fluorescent markers were singled and successful integration of GFP-SEC was confirmed by PCR spanning the homology regions on both sides. Roller worms (*gfp-SEC::zen-4/+*) were mated with the *nT1[qIs51]* balancer to facilitate SEC removal. The L1/L2 larvae of balanced roller worms (*gfp-SEC::zen-4/nT1[qIs51])* were heat-shocked at 34°C for 4 hours to induce Cre expression and SEC removal. Normal-moving progeny without the *nT1[qIs51]* balancer (homozygous *gfp::zen-4*) were selected. Successful SEC excision and correct GFP integration were confirmed by PCR across both sides of the GFP insertion.

### RNA interference

Single-stranded RNAs (ssRNAs) were synthesized in 50 μL T3 and T7 reactions (MEGAscript, Invitrogen, Carlsbad, CA) using cleaned DNA templates generated by PCR from N2 genomic DNA or cDNA using oligonucleotides containing T3 or T7 promoters (Table S2). Reactions were cleaned using the MEGAclear kit (Invitrogen, Carlsbad, CA), and the 50 μL T3 and T7 reactions were mixed with 50 μL of 3x soaking buffer (32.7 mM Na_2_HPO_4_, 16.5 mM KH_2_PO_4_, 6.3 mM NaCl, 14.1 mM NH_4_Cl) and annealed (68°C for 10 min followed by 37°C for 30 min). L4 hermaphrodites were injected with dsRNA and incubated at 16°C or 20°C depending on the experiment. For double or triple depletions, dsRNAs were mixed at equal concentrations (∼2 μg/μl for each dsRNA); a dsRNA targeting *hyls-1*, a gene with no known function in the early embryo, was used as a mixing control.

To count number of embryos laid after RNAi-mediated depletion, L4 hermaphrodites were injected with dsRNAs and incubated at 20°C for 24 hours (or as indicated in specific experiments). Worms were singled and allowed to lay embryos at 20°C for 24 hours (or as indicated in specific experiments), adult worms were removed and all embryos and hatchlings were counted.

For the imaging experiment in Figure 4C, the strains used also contained a *cyk-*4-RNAi-resistant transgene encoding CYK-4::GFP::MEI-1(1–224) that was originally engineered for experiments not presented in this study. L4 hermaphrodites were co-injected with *cyk-4* dsRNA (to deplete endogenous CYK-4) and *gfp* dsRNA (to deplete CYK-4::GFP::MEI-1(1–224)) and incubated at 20°C for 48 hours before imaging.

### Imaging experiments

Germline imaging was performed by anesthetizing worms in 1 mg/ml Tricane (ethyl 3-aminobenzoate methanesulfonate salt) and 0.1 mg/ml of tetramisole hydrochloride (TMHC) dissolved in M9 for 15–30 minutes. Anesthetized worms were transferred to a 2% agarose pad, overlaid with coverslip, and imaged using a spinning disk confocal system (Andor Revolution XD Confocal System; Andor Technology) with a confocal scanner unit (CSU-10; Yokogawa) mounted on an inverted microscope (TE2000-E; Nikon) equipped with a 60×/1.4 Plan-Apochromat objective, solid-state 100-mW lasers, and an electron multiplication back-thinned charge-coupled device camera (iXon; Andor Technology. Germline imaging was performed by acquiring a 40 x 1 μm z-series, with no binning (Figures 1C-D, 2A, 2C, 2E, 4C, 4D, 5B-D and Figure 1—figure supplement 4A), or with 2x2 binning (Figures 2D, 5B). For Figure 1B, Figure 1—figure supplement 1, and Figure 1—figure supplement 2, 61 x 0.5 μm (L1–L4 germlines) or 81 x 0.5 μm z-series (adult germlines) were acquired. For Figure 2-figure supplement 2, a 200 x0.2 μm z-series was collected and for Figure 5-figure supplement 1, a 80 x 0.5 μm z-series was collected. For compartment bridge diameter measurement in Figure 1—figure supplement 3, young adult germlines were imaged by acquiring a 129 x 0.25 μm z-series with no binning. Cross sections were used to measure the length of the CYK-4::mNeonGreen signal across the widest opening of each compartment bridge.

### Construction of worm trap microfluidic devices

The worm trap microfluidic device (**Figure 3—figure supplement 1**) is assembled out of a 7 mm thick polydimethylsiloxane (PDMS) chip with microchannels engraved on its surface and a 35x50 mm #1.5 microscope coverslip, which seals the microchannels. The microchannels of the device are of three different depths, 10, 50, and 750 μm. PDMS chips were cast out of a 50/50 mixture of Sylgard 184 (Dow-Corning) and XP-592 (Silicones Inc.) silicone pre-polymers, each of which was a 10:1 mixture of the base and cross-linker components of the respective silicone material. The master mold used to cast the chips was a 5-inch silicon wafer with a photolithographically fabricated microstructure with 10, 50, and 750 μm tall features made of UV-cross-linked photoresists of the SU8 family (MicroChem, Newton, MA). To fabricate the master mold, the wafer was spin-coated with SU8 2005 to a thickness of 10 μm, baked and exposed to UV light through a photomask. After that, the wafer was spin-coated with SU8 2015 to a cumulative thickness of 50 μm, exposed through another photomask, and baked. Finally, the wafer was spin-coated with SU8 2150 to a cumulative thickness of 750 μm, exposed through a third photomask, baked, and developed.

To facilitate the immobilization of worms in the imaging chambers, the surfaces of both PDMS (ceiling of the imaging chambers) and glass coverslip (floor of the imaging chambers) were roughened by coating them with 0.2 μm polystyrene beads, which were covalently attached to the surfaces using carboxyl-amine chemistry. (The size of the beads was too small to cause substantial scattering of light, making the beads practically invisible under a microscope.) To this end, the surfaces of both the coverslip and PDMS chip were oxidized by exposing them for 10 seconds to oxygen plasma (using a PlasmaPreen II plasma treater by Plastmatic Systems Inc.) and immediately treated with a solution aminopropyltrimethoxysilane (4% in EtOH) and then incubated under this solution for 5 minutes at room temperature (in a fume hood). As a result of this treatment, the glass and PDMS surfaces were coated with amine groups. The coverslip and PDMS chip were rinsed with ethanol, dried with filtered compressed air, and then incubated for 30 min at room temperature under a 0.01% suspension of 0.20 μm carboxylated polystyrene beads (Polybead^^®^^ Carboxylate Microspheres, from Polysciences) in a 20 mM solution of HEPES, pH 8, containing 0.01% EDC [1-Ethyl-3-(3-dimethylaminopropyl)carbodiimide; to promote the reaction between carboxyl groups on the beads and amine groups on the surface of glass or PDMS]. The coverslip and PDMS chip were then rinsed with water and dried.

### Experiments in worm trap microfluidic device

To load *cyk-4* mutant animals (OD239 *(cyk-4 (or749ts))* and OD241 *(cyk-4(t1689ts))*) into the worm trap device, the PDMS chip was placed with its microchannels facing up, and a 1ul droplet of media (0.5xPBS + 1:10 dilution of HB101 overnight culture) was dispensed onto the chip near the center of each of the three 50μm deep, 3.7x3 mm rectangular imaging chambers on the chip surface (the drops remained separate because of hydrophobicity of PDMS). One or more young adult worms were carefully transferred into each drop using a platinum pick. The chip was inverted, and the engraved bottom side was brought in contact with the coverslip; this turned the microgrooves into sealed microchannels and the drop medium filled these microchannels. To hold the chip and coverslip together, a deep O-shaped channel, which surrounded the liquid-filled microchannels of the device and served as a vacuum cup,was connected to a regulated source of vacuum that was set at a low level of -5 kPa in the gauge pressure (low vacuum). The application of vacuum resulted in a partial collapse of the imaging chambers, reducing their depth to ∼45 μm, which did not prevent worms from moving freely and feeding. To trap worms for high-resolution imaging, the adjustable vacuum level was dialed up to -25 kPa (high vacuum), causing a major reduction of the imaging chamber depths and immediate immobilization of worms in the chambers. If immobilized worms were poorly positioned, the vacuum was dialed back to low, allowing worms to move, and then to high again, re-trapping worms for imaging. The immobilization was most effective away from the edges of the imaging chambers. Imaging was performed on a spinning disk confocal microscope (Andor Revolution XD Confocal System; Andor Technology) as described above. A stack of 40 images with a 1 μm step along the Z-axis was collected. After imaging was completed, the vacuum was switched from high to low, allowing worms to move, feed, and recover until next imaging timepoint.

Initial stacks of confocal images in the temperature shift experiments were acquired immediately after the device was assembled, without perfusion of a bacterial culture through the device. At temperature shift, the device inlet was connected through a tubing line to a reservoir with bacterial culture (which was immersed in a 38°C water bath), the device outlet was connected to a reservoir with plain medium, and a continuous perfusion of bacterial culture through the imaging chambers was initiated by setting a constant positive differential pressure between the inlet and outlet (by lifting the inlet reservoir above the outlet reservoir). In preparation for temperature shift to 25°C, a warm air blower was positioned adjacent to the stage, with its air flow directed towards the worm trap device and the tubing line connected to the device inlet. A temperature probe was affixed to the stage adjacent to the device to monitor the temperature throughout the course of the experiment; the blower was manually adjusted to maintain air temperature on the stage at 25.5-26.5 °C during the course of the experiment.

Prior to trapping and imaging at the initial restrictive time point, worms were exposed to the restrictive temperature for ∼45 minutes. Subsequently, worms were trapped and imaged every 45-60min over a period of 5 hours. For most experiments, two worm trap devices were used in parallel to enable side-by-side analysis of the two *cyk-4* mutants (*t1689ts,* and *or749ts*). Each device was operated independently of the other with respect to both perfusion and the application of vacuum. (Two separate vacuum regulators set at -5 and -25 kPa made it possible to independently apply either low or high vacuum to either device.) Thus, initial perfusion, trapping and imaging was offset by 20-30 min and subsequent imaging alternated between each of the two traps every 20-30 minutes during the time course. Three independent experiments were performed for each strain (n=7 for *cyk-4(t1689ts),* and n=9 for *cyk-4(or749ts)*).

### Image analysis

All images were processed, scaled, and analyzed using ImageJ (National Institutes of Health) or MetaMorph software (Molecular Devices).

### Western blotting

To reduce the contamination with *E. coli* (which the worms eat), 45 worms from each of the indicated conditions were transferred onto a medium plate with no food and the plate was flooded with 4 ml of M9. After 1 hour, when the *E. coli* have largely been flushed out of the worms’ digestive tracts, the worms were transferred into a tube containing 500 μl of M9 in a screw-cap 1.5 ml tube, 0.5 ml of M9 + 0.1% Triton X-100 was added, and worms were pelleted by centrifuging at 400 xg for 1–2 min. Buffer was removed leaving behind 50 μl and worms were washed and pelleted three more times in 1 ml of M9 + 0.1% Triton X-100. After the last wash, buffer was removed leaving behind 30 μl and 10 μl of 4X sample buffer was added. Samples were placed in a sonicating water bath at 70°C for 10 min, boiled in a 95°C heating block for 5 min, and sonicated again for 10 min at 70°C before freezing. Samples were thawed and separated by SDS-PAGE. After transferring the separated protein bands to a nitrocellulose membrane, the blot was cut into two parts at ∼70 kDa. The above-70-kDa blot was probed using 1 μg/ml of rabbit anti-CYK-4 (aa 131–345), which was detected using an HRP-conjugated secondary antibody (1:10,000; GE Healthcare Life Sciences) and WesternBright Sirius detection system (Advansta). The below-70-kDa blot was probed for α-tubulin using the monoclonal DM1α antibody (1:500; Sigma-Aldrich), and then visualized by Western Blue Stabilized Substrate for Alkaline Phosphatase (1:2,000; Promega). Antibodies against CYK-4 were generated by injecting a purified GST fusion with CYK-4 aa 131–345 into rabbits (Covance). Antibodies were purified from serum using standard procedures (Harlow, 1988) on a 1 ml NHS HiTrap column (GE Healthcare Life Sciences) containing an immobilized MBP fusion with the same region of CYK-4.

### Positional correlative anatomy EM

Samples were high pressure frozen followed by freeze-substitution as described previously (Kolotuev, Schwab, & Labouesse, 2009) and flat embedded, targeted and sectioned using the positional correlation and tight trimming approach (Kolotuev, 2014). For ultramicrotome sectioning, a Leica UC7 microtome was used. 100nm sections were collected on the slot-formvar coated grids and observed using a JEOL JEM 1400 TEM microscope (JEOL, Japan). Samples were aligned and rendered using the ImageJ and IMOD programs.

## ACKNOWLEDGEMENTS

We would like to thank the electron microscopy facilities in Rennes and Lausanne, and Tom Garber for assistance in constructing the EM model. J.S.G-C was supported by the University of California, San Diego Cancer Cell Biology Training Program (T32 CA067754). A.D. and K.O. receive salary and other support from the Ludwig Institute for Cancer Research. A.G and E.G. were supported by an award from the National Science Foundation (PHY-14113130 to A.G.).

---

**Figure 1—figure supplement 1.**
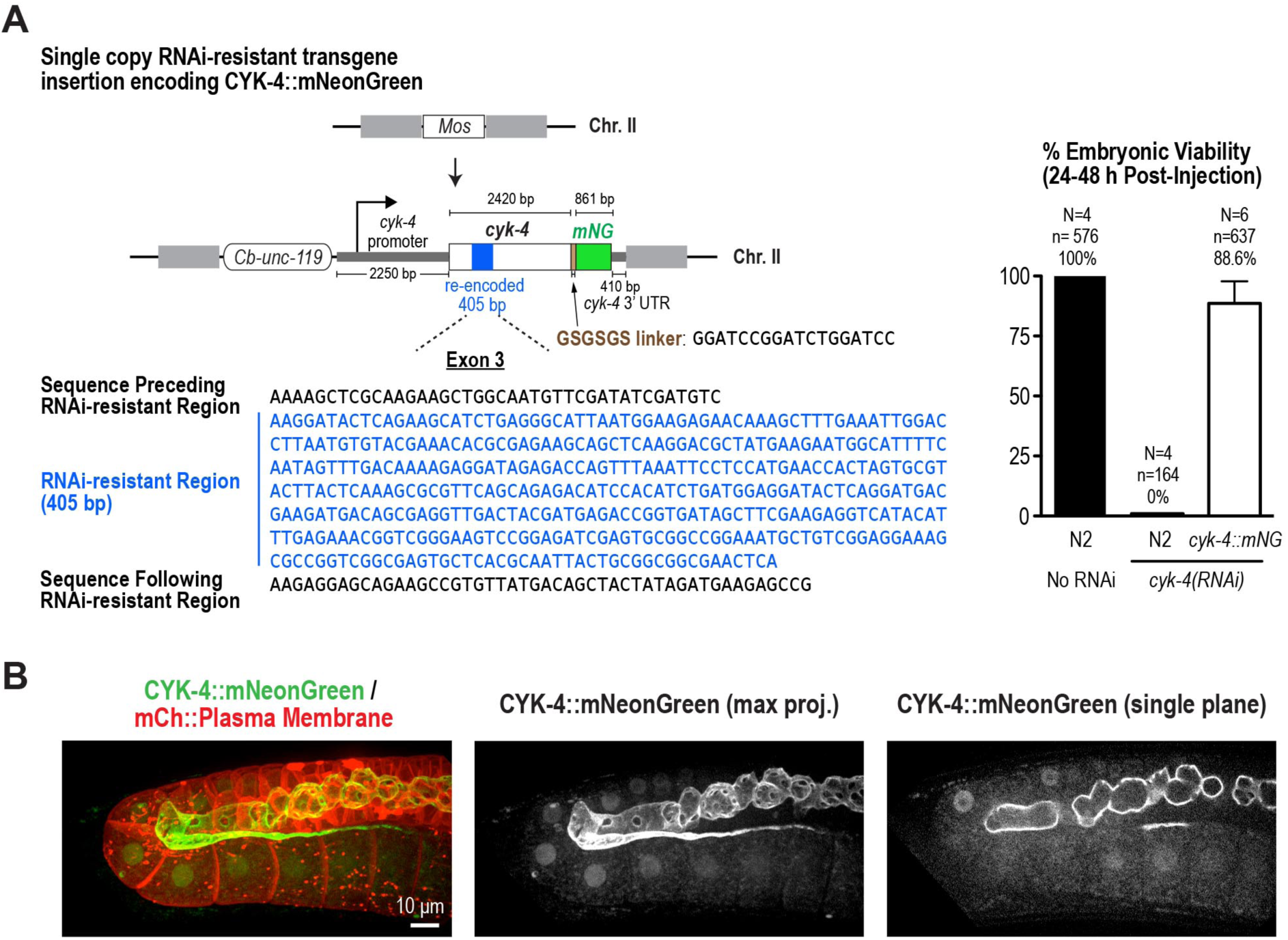
Generation of a functional single-copy transgene encoding CYK-4::mNeonGreen. (**A**) (*Left*) Schematic of the RNAi-resistant *cyk-4::mNeonGreen* transgene integrated into a specific site (Mos transposon insertion) on Chromosome II. (*Right*) Graph plotting embryonic viability (mean ± SD) following depletion of endogenous CYK-4 by RNAi. N = number of worms, n = number of embryos. (**B**) Fluorescence confocal images of an adult germline in a worm expressing CYK-4::mNeonGreen and an mCherry-tagged plasma membrane probe. The images below the merge show a maximum intensity projection (merged imaged reproduced from Figure 1B for comparison with the single color image) and a single central plane image for the CYK-4::mNeonGreen channel. Scale bar, 10 μm.

**Figure 1—figure supplement 2.**
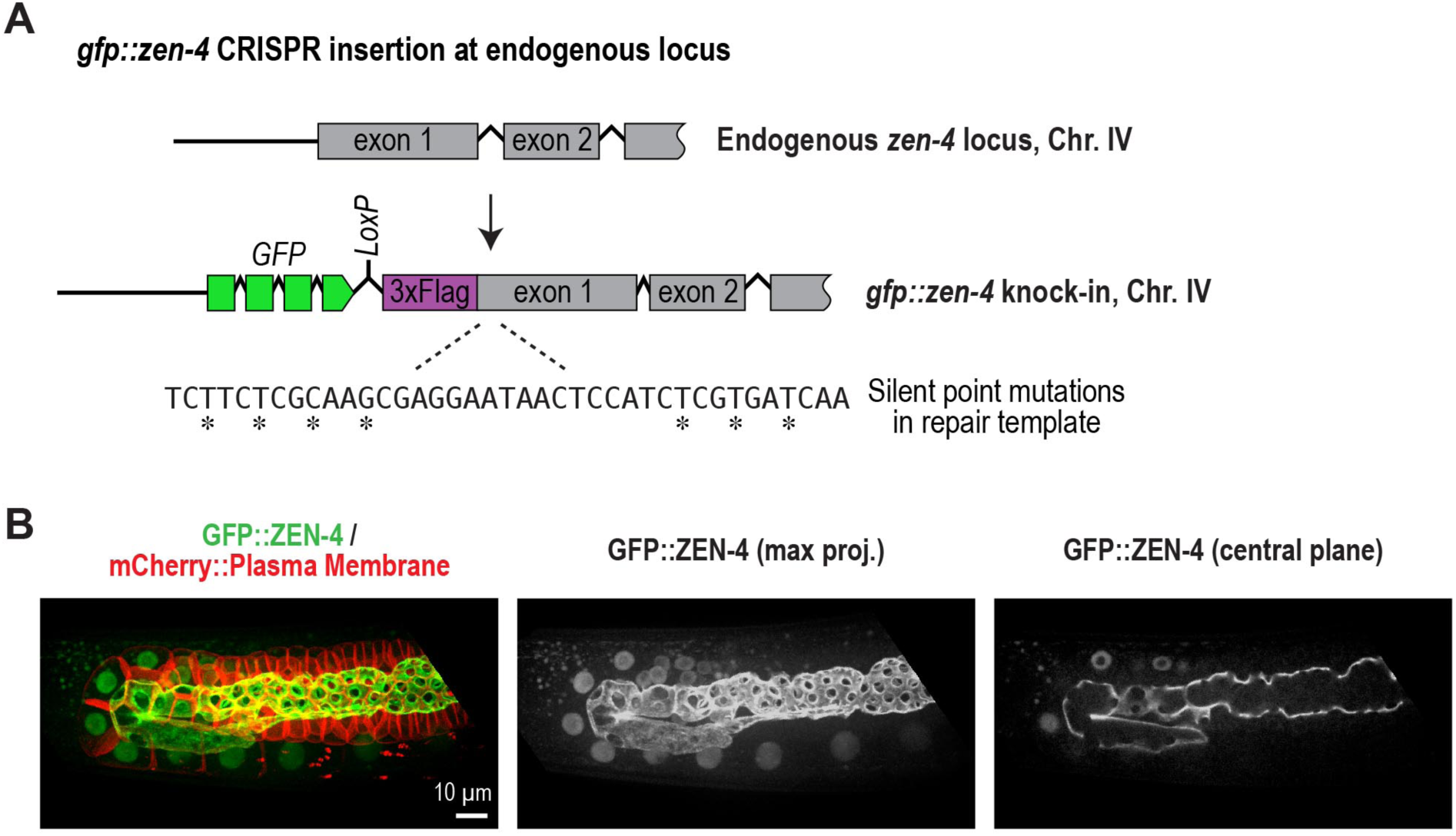
Generation of *in situ* tagged GFP::ZEN-4. (**A**) Schematic showing the location where sequences encoding GFP were inserted prior to the first exon of the gene encoding ZEN-4. (**B**) Fluorescence confocal images of an adult germline in a worm expressing GFP::ZEN-4 and an mCherry-tagged plasma membrane probe. The images below the merge show a maximum intensity projection (merged imaged reproduced from Figure 1B for comparison with the single color image) and a single central plane image for the GFP::ZEN-4 channel. Scale bar, 10 μm.

**Figure 1—figure supplement 3.**
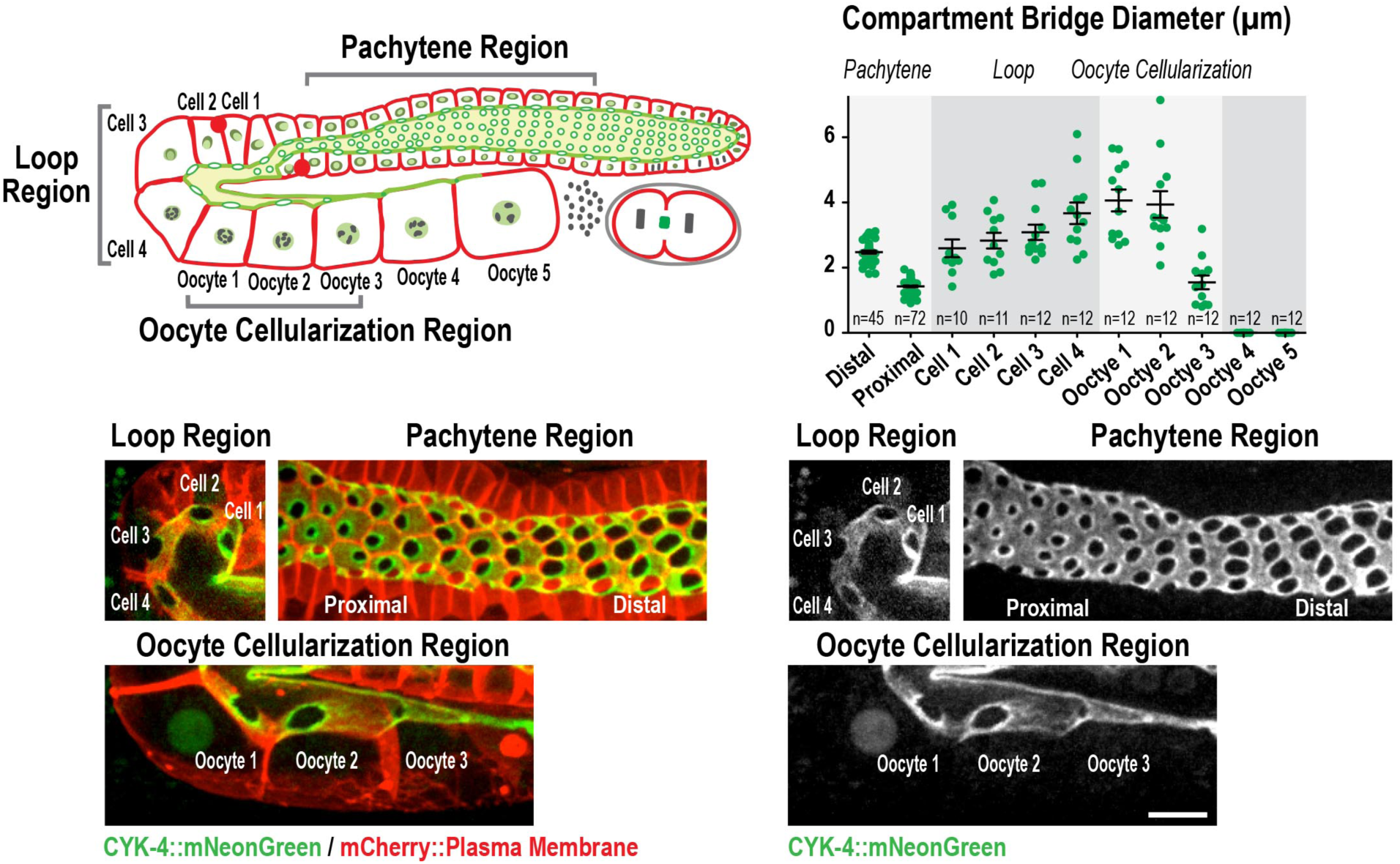
Compartment bridges expand during oocyte loading and close during cellularization. (*Upper left*) Schematic shows the location of the germline regions in the images and the nomenclature for compartment and oocyte labeling. (*Upper right*) Graph plotting the diameters of the compartment bridges in the indicated regions of the germline measured in the CYK-4::mNeonGreen images. n = number of cells analyzed at the indicated position/region. (*Lower panels*) Maximum intensity projections of confocal images of the pachytene, loop, and oocyte cellularization regions of adult germlines acquired in the strain expressing an mCherry-tagged plasma membrane marker and CYK-4::mNeonGreen. Merged images (reproduced from Figure 1B) are shown alongside single color images showing the CYK-4::mNeonGreen signal. Scale bar, 10 μm.

**Figure 1—figure supplement 4.**
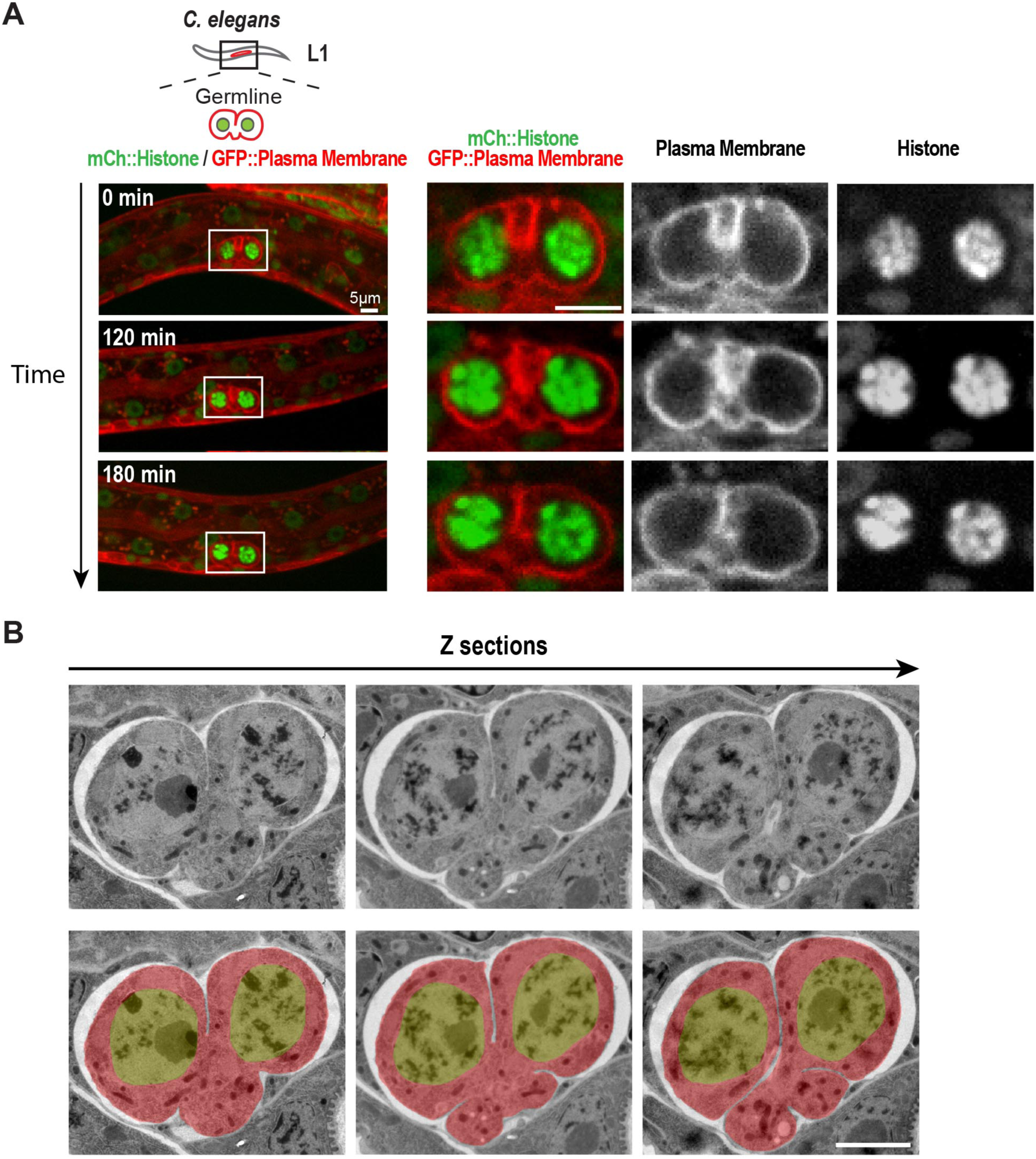
Structure of the nascent syncytial germline and rachis in the L1 larva. (**A**) Timelapse images collected using the worm trap described in Figure 3A over a 3-hour period of the germline in an L1 stage worm. At this stage, the intercellular bridge connecting the two nuclear compartments extends out on one side. Scale bars are 5 μm. (**B**) Selected images from a stack of serial 100 nm sections of an L1 germline (to view the entire image series see Video 1) that were collected and imaged by transmission electron microscopy after fixation by high pressure freezing and freeze-substitution. Images are shown without *(top*) and with (*bottom*) superposition of a pseudocolor model in which the regions within the boundary of the 2-compartment germline syncytium (*red*) and germline nuclei (*green*) are highlighted. The two nuclei-containing compartments sit side-by-side and open into a small cytoplasm containing bridge (nascent rachis) that extends out to one side. Images above and below the central planes reveal that the intercellular bridge/rachis has a lobed structure. Scale bar is 2 μm.

**Figure 1—figure supplement 5.**
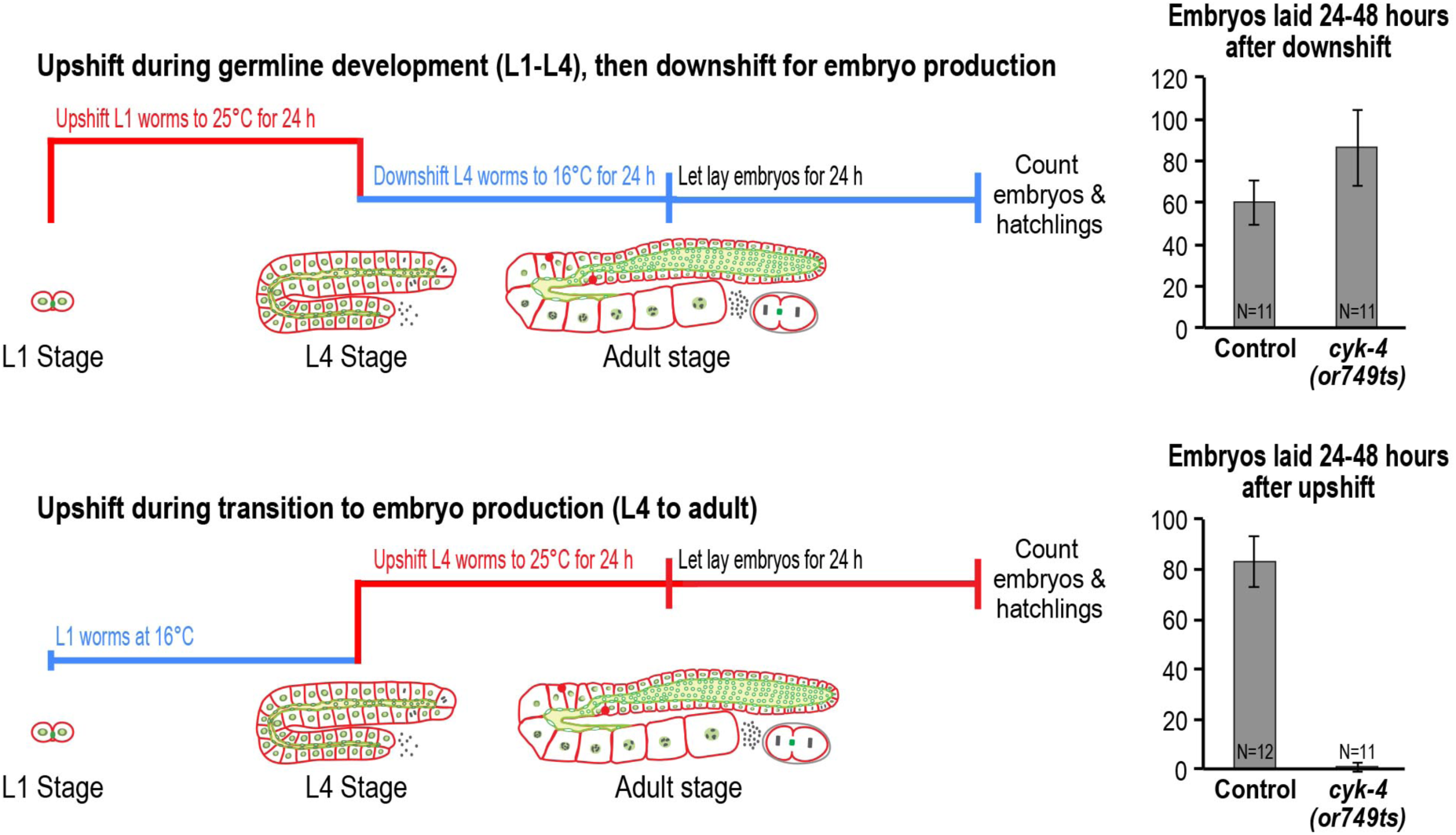
CYK-4 is required for embryo production by the adult syncytial germline. (*Left*) Schematic outlines of the upshift protocols. (*Right*) Graphs plot the number of embryos laid during the indicated intervals. N= number of worms.

**Figure 2—figure supplement 1.**
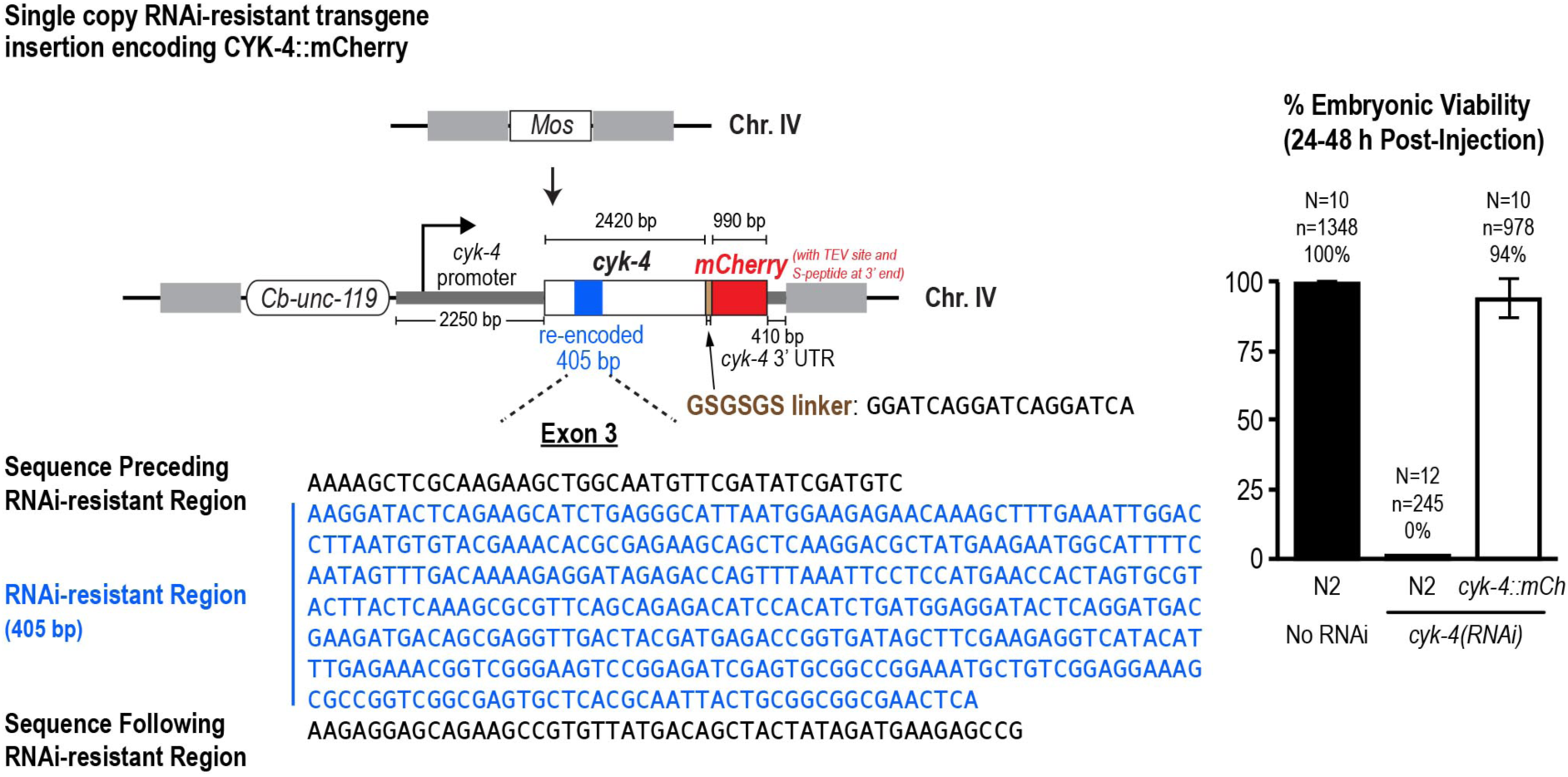
Generation of a functional single-copy transgene encoding CYK-4::mCherry. (**A**) (*Left*) Schematic of the RNAi-resistant *cyk-4::mCherry* transgene integrated into a specific site (Mos transposon insertion) on Chromosome IV. (*Right*) Graph plotting embryonic viability (mean ± SD) following depletion of endogenous CYK-4 by RNAi. N = number of worms, n = number of embryos.

**Figure 2—figure supplement 2.**
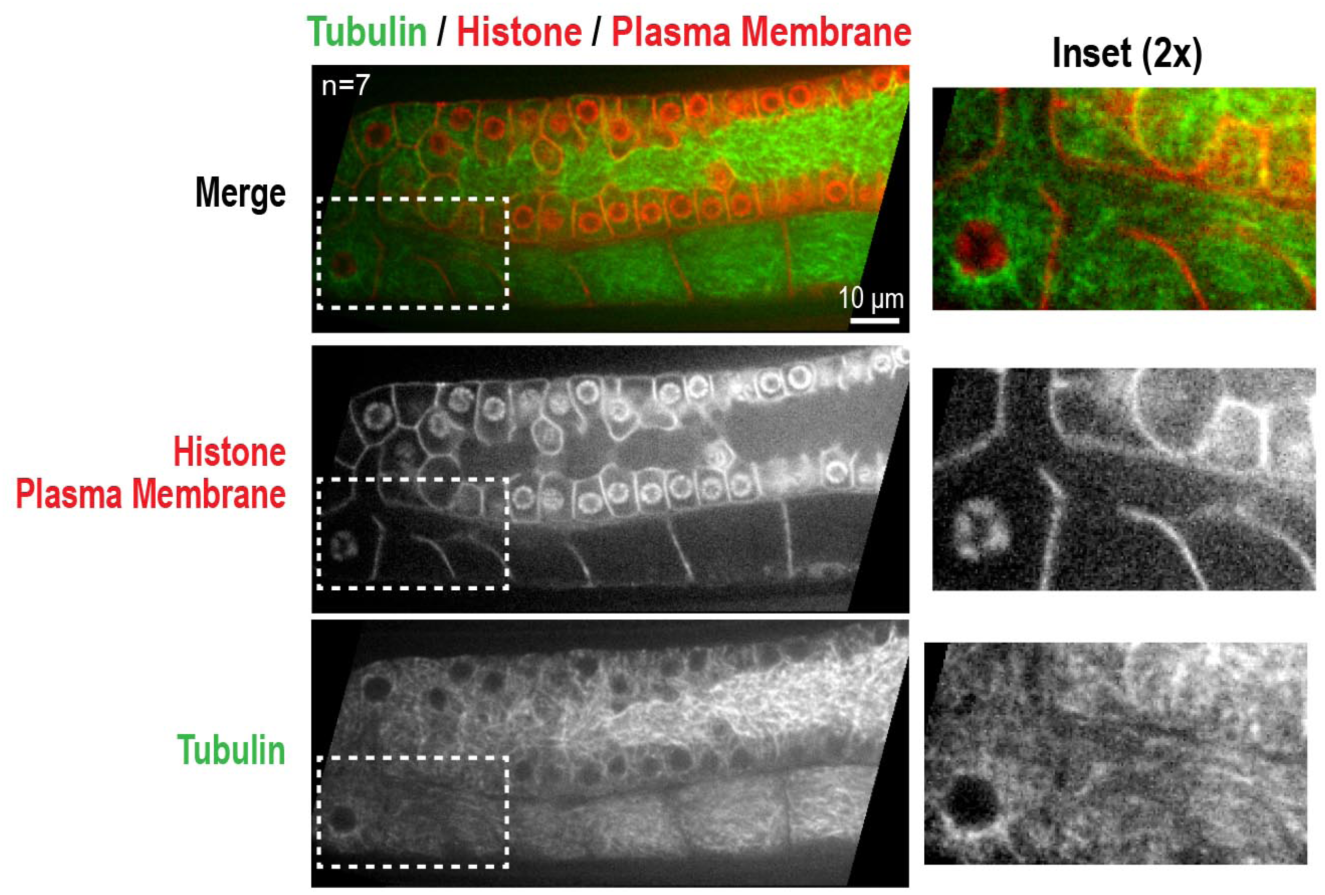
The microtubule cytoskeleton in the pachytene, loop, and oocyte cellularization region of the germline. (*Top*) Single plane confocal image of an adult germline in a worm expressing GFP::β-tubulin, mCherry::histone and an mCherry-tagged plasma membrane probe (reproduced from Figure 2F for comparison). The images below the merge show the red and green channels separately. Scale bar, 10 μm.

**Figure 3—Figure Supplement 1.**
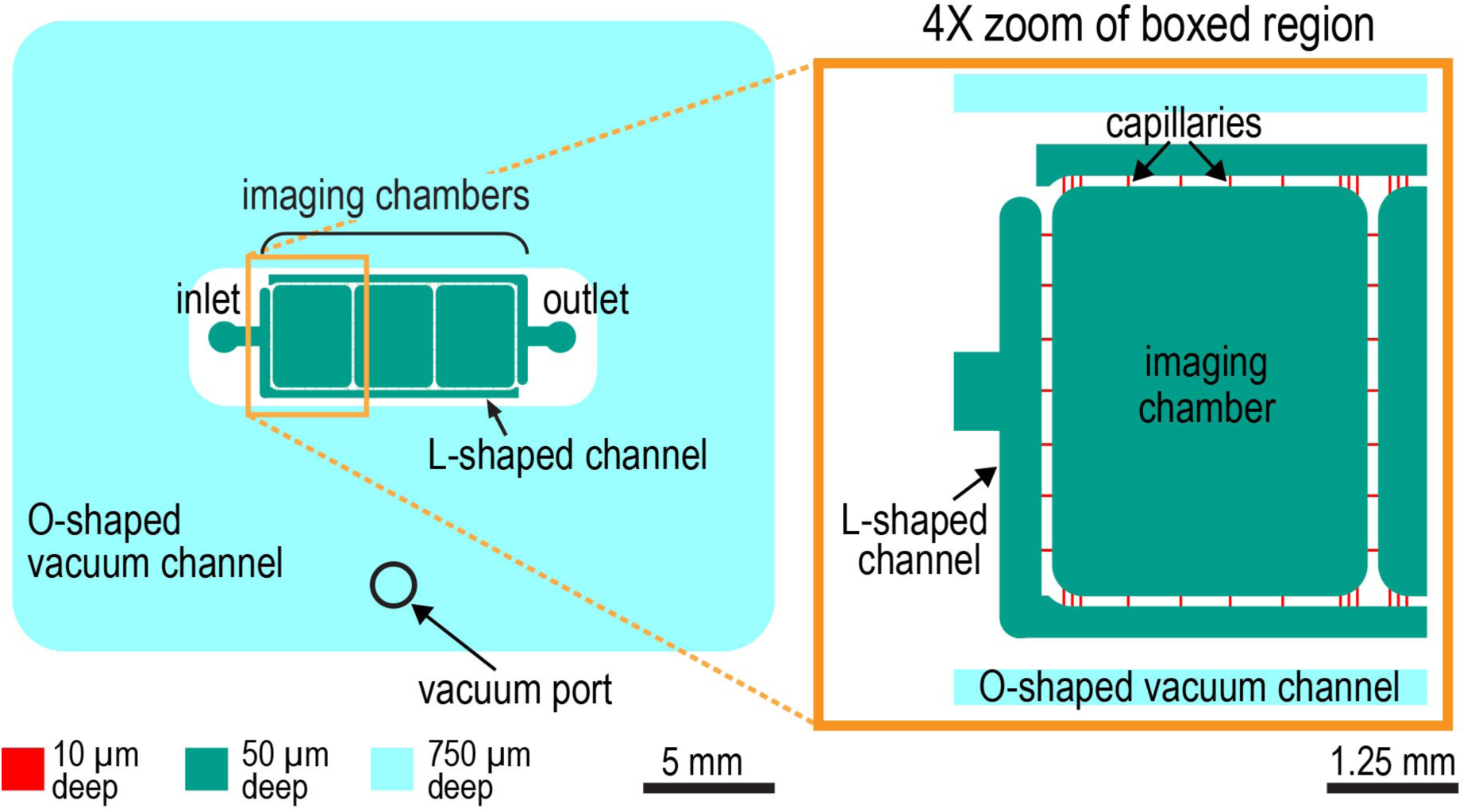
Design of the worm trap microfluidic chip. Schematic of microfluidic device (*left*) and 4X zoom of left-most imaging chamber (*right*). The worm trap microfluidic device is assembled out of a PDMS chip with microchannels engraved on its surface and a 35x50 mm #1.5 microscope coverslip, which seals the microchannels (*not shown*). The microchannels of the device are of three different depths, 10, 50, and 750 μm (shown in red, green, and cyan, respectively). The 10 and 50 μm deep microchannels form a liquid-filled network with one inlet and one outlet as indicated that is surrounded by a separate O-shaped 750 μm deep channel (*cyan*). This last channel serves as a vacuum cup. When the chip is sealed with a coverslip, and vacuum is applied to this 750 μm deep channel (through a dedicated vacuum port) the chip and coverslip are pulled together. The application of vacuum also leads to partial collapse of all channels of the device, including the liquid-filled microchannels, making it possible to dynamically reduce the microchannel depth in a controlled way by adjusting the level of vacuum. The main functional elements of the liquid-filled microchannel network are the three identical 3.7x3 mm imaging chambers, one of which is highlighted in zoom panel, which are 50 μm deep and have rounded corners. The 50 μm depth (as well as somewhat reduced depths of 40-45 μm, when the chambers partially collapse under a low vacuum of -5kPa) was empirically found to be sufficient for larval and young adult worms to move freely and feed. The imaging chambers are connected to each other and to the device inlet and outlet through capillary microchannels with cross-sections of 10x10 μm. The small cross-section of these capillary microchannels makes it impossible for worms to enter them. Hence, worms cannot escape from the imaging chambers. The inlet and outlet are connected to the capillary microchannels through 50 μm deep, 300 μm wide L-shaped microchannels. Relatively low fluidic resistance of these last microchannels (as compared with the 10x10 μm capillaries) facilitates even distribution of flow between the capillaries and even perfusion of the imaging chambers with bacterial suspension, which introduced via the inlet.

**Figure 4—figure supplement 1.**
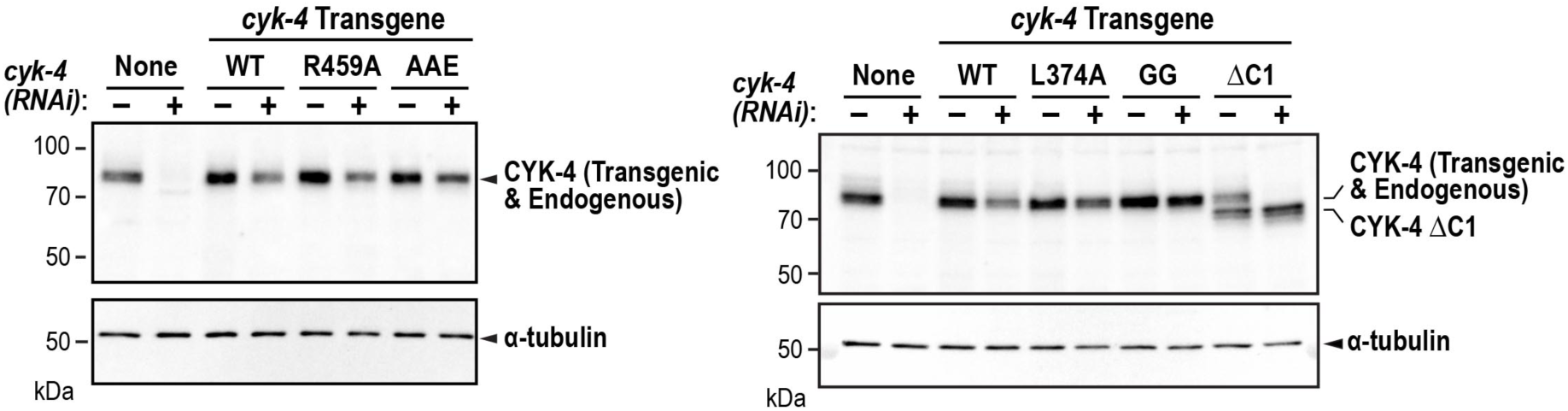
The WT, GTPase binding interface mutant, and ΔC1 mutant proteins encoded by the single copy untagged transgenes are expressed at levels comparable to endogenous CYK-4. Full versions of the blots in Figure 4B. L374A and GG are point mutants in the C1 domain that are not used in this study.

**Figure 5—figure supplement 1.**
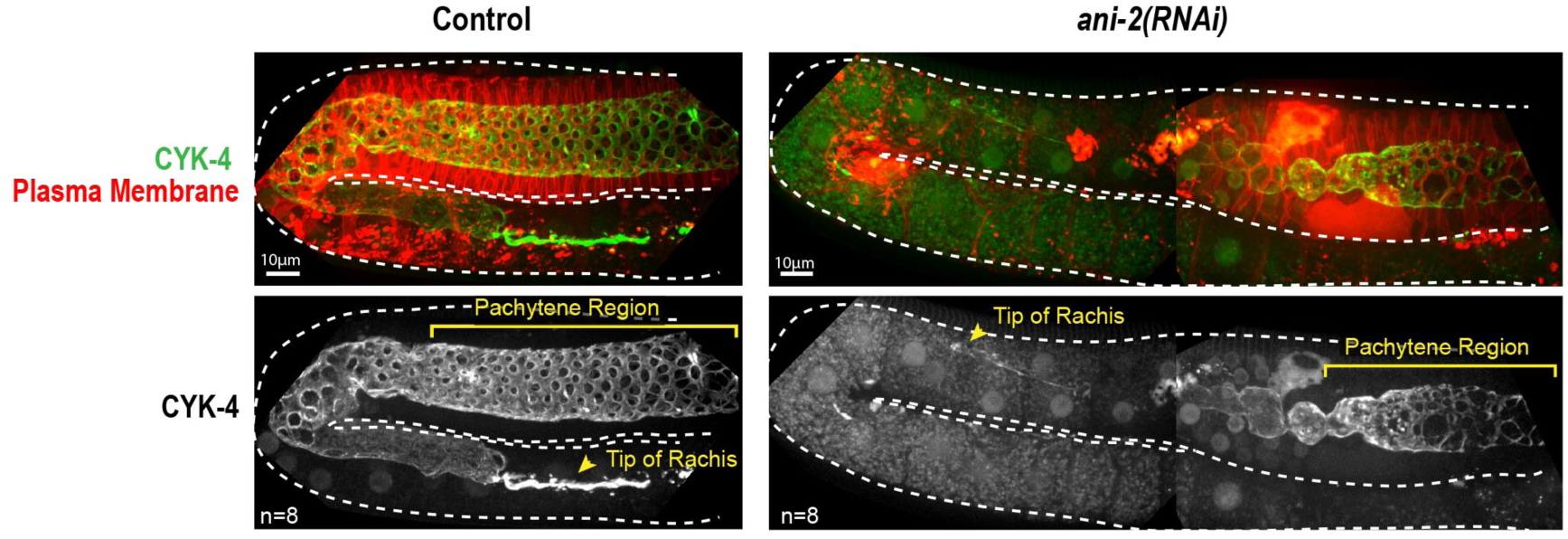
The anillin homolog, ANI-2, is not required for CYK-4 recruitment to the rachis surface or compartment bridges. Maximum intensity projections of germlines in adult control and *ani-2(RNAi)* (48 hours after introduction of dsRNA by soaking) animals expressing CYK-4::mNeonGreen and an mCherry tagged plasma membrane probe. Dotted region outlines the boundary of the germline. ANI-2 depletion leads to germline defects that increase the number of unfertilized oocytes within the germline and push the rachis tip and pachytene region backwards within the body of the worm. Thus, a larger region of *ani-2(RNAi)* worms was imaged and the images were stitched together to allow comparison of the ability of CYK-4 to localize to the rachis surface. n = number of worms. Scale bar is 10 μm.

**Figure 5—figure supplement 2.**
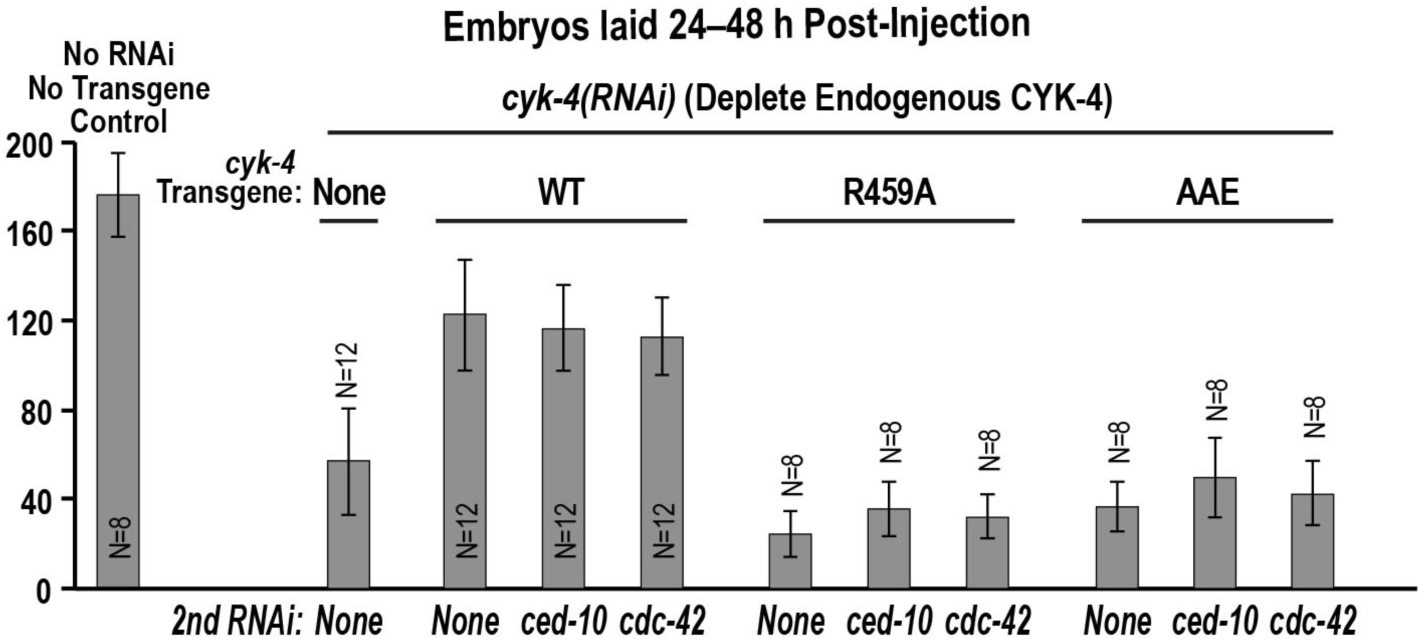
Depletion of Rac^CED-10^ or CDC-42 cannot rescue the effects of mutants in the Rho GTPase binding interface on the germline. Graph shows the number of eggs laid 24 – 48 hours post-injection (mean ± SD) for the indicated strains and RNAi conditions. N = number of worms.

**VIDEO LEGEND**

**Video 1: Structure of the syncytial germline at the 2-compartment stage in an L1 larva.** Video shows images of a stack of serial 100 nm serial sections of an L1 germline that were collected and imaged by transmission electron microscopy. Images are shown without and then with superposition of a pseudocolor model in which the regions within the boundary of the 2-compartment germline syncytium (*red*) and germline nuclei (*green*) are highlighted.

## REFERENCES

Agromayor, M., & Martin-Serrano, J. (2013). Knowing when to cut and run: mechanisms that control cytokinetic abscission. Trends Cell Biol, 23(9), 433–441. doi:10.1016/j.tcb.2013.04.006

Amini, R., Goupil, E., Labella, S., Zetka, M., Maddox, A. S., Labbe, J. C., & Chartier, N. T. (2014). C. elegans Anillin proteins regulate intercellular bridge stability and germline syncytial organization. J Cell Biol, 206(1), 129–143. doi:10.1083/jcb.201310117

Arur, S., Ohmachi, M., Nayak, S., Hayes, M., Miranda, A., Hay, A., … Schedl, T. (2009). Multiple ERK substrates execute single biological processes in Caenorhabditis elegans germ-line development. Proc Natl Acad Sci U S A, 106(12), 4776–4781. doi:10.1073/pnas.0812285106

Basant, A., Lekomtsev, S., Tse, Y. C., Zhang, D., Longhini, K. M., Petronczki, M., & Glotzer, M. (2015). Aurora B kinase promotes cytokinesis by inducing centralspindlin oligomers that associate with the plasma membrane. Dev Cell, 33(2), 204–215. doi:10.1016/j.devcel.2015.03.015

Breznau, E. B., Murt, M., Blasius, T. L., Verhey, K. J., & Miller, A. L. (2017). The MgcRacGAP SxIP motif tethers Centralspindlin to microtubule plus ends in Xenopus laevis. J Cell Sci, 130(10), 1809–1821. doi:10.1242/jcs.195891

Burkard, M. E., Maciejowski, J., Rodriguez-Bravo, V., Repka, M., Lowery, D. M., Clauser, K. R., … Jallepalli, P. V. (2009). Plk1 self-organization and priming phosphorylation of HsCYK-4 at the spindle midzone regulate the onset of division in human cells. PLoS Biol, 7(5), e1000111. doi:10.1371/journal.pbio.1000111

Canman, J. C., Lewellyn, L., Laband, K., Smerdon, S. J., Desai, A., Bowerman, B., & Oegema, K. (2008). Inhibition of Rac by the GAP activity of centralspindlin is essential for cytokinesis. Science, 322(5907), 1543–1546. doi:10.1126/science.1163086

Carmena, M., Riparbelli, M. G., Minestrini, G., Tavares, A. M., Adams, R., Callaini, G., & Glover, D. M. (1998). Drosophila polo kinase is required for cytokinesis. J Cell Biol, 143(3), 659–671.

Davies, T., Jordan, S. N., Chand, V., Sees, J. A., Laband, K., Carvalho, A. X., … Canman, J. C. (2014). High-resolution temporal analysis reveals a functional timeline for the molecular regulation of cytokinesis. Dev Cell, 30(2), 209–223. doi:10.1016/j.devcel.2014.05.009

Dickinson, D. J., Pani, A. M., Heppert, J. K., Higgins, C. D., & Goldstein, B. (2015). Streamlined Genome Engineering with a Self-Excising Drug Selection Cassette. Genetics, 200(4), 1035–1049. doi:10.1534/genetics.115.178335

Dym, M., & Fawcett, D. W. (1971). Further observations on the numbers of spermatogonia, spermatocytes, and spermatids connected by intercellular bridges in the mammalian testis. Biol Reprod, 4(2), 195–215.

Elia, N., Sougrat, R., Spurlin, T. A., Hurley, J. H., & Lippincott-Schwartz, J. (2011). Dynamics of endosomal sorting complex required for transport (ESCRT) machinery during cytokinesis and its role in abscission. Proc Natl Acad Sci U S A, 108(12), 4846–4851. doi:10.1073/pnas.1102714108

Encalada, S. E., Martin, P. R., Phillips, J. B., Lyczak, R., Hamill, D. R., Swan, K. A., & Bowerman, B. (2000). DNA replication defects delay cell division and disrupt cell polarity in early Caenorhabditis elegans embryos. Dev Biol, 228(2), 225–238. doi:10.1006/dbio.2000.9965

Foe, V. E., & von Dassow, G. (2008). Stable and dynamic microtubules coordinately shape the myosin activation zone during cytokinetic furrow formation. J Cell Biol, 183(3), 457–470. doi:10.1083/jcb.200807128

Frokjaer-Jensen, C., Davis, M. W., Hopkins, C. E., Newman, B. J., Thummel, J. M., Olesen, S. P., … Jorgensen, E. M. (2008). Single-copy insertion of transgenes in Caenorhabditis elegans. Nat Genet, 40(11), 1375–1383. doi:10.1038/ng.248

Gibert, M. A., Starck, J., & Beguet, B. (1984). Role of the gonad cytoplasmic core during oogenesis of the nematode Caenorhabditis elegans. Biol Cell, 50(1), 77–85.

Gonczy, P., Schnabel, H., Kaletta, T., Amores, A. D., Hyman, T., & Schnabel, R. (1999). Dissection of cell division processes in the one cell stage Caenorhabditis elegans embryo by mutational analysis. J Cell Biol, 144(5), 927–946.

Goupil, E., Amini, R., & Labbe, J.-C. (2017). Anillin proteins stabilize the cytoplasmic bridge between the two primordial germ cells during C. elegans embryogenesis. bioRxiv. doi:10.1101/126284

Green, R. A., Kao, H. L., Audhya, A., Arur, S., Mayers, J. R., Fridolfsson, H. N., … Oegema, K. (2011). A high-resolution C. elegans essential gene network based on phenotypic profiling of a complex tissue. Cell, 145(3), 470–482. doi:10.1016/j.cell.2011.03.037

Green, R. A., Mayers, J. R., Wang, S., Lewellyn, L., Desai, A., Audhya, A., & Oegema, K. (2013). The midbody ring scaffolds the abscission machinery in the absence of midbody microtubules. J Cell Biol, 203(3), 505–520. doi:10.1083/jcb.201306036

Green, R. A., Paluch, E., & Oegema, K. (2012). Cytokinesis in animal cells. Annu Rev Cell Dev Biol, 28, 29–58. doi:10.1146/annurev-cellbio-101011-155718

Greenbaum, M. P., Iwamori, N., Agno, J. E., & Matzuk, M. M. (2009). Mouse TEX14 is required for embryonic germ cell intercellular bridges but not female fertility. Biol Reprod, 80(3), 449–457. doi:10.1095/biolreprod.108.070649

Greenbaum, M. P., Ma, L., & Matzuk, M. M. (2007). Conversion of midbodies into germ cell intercellular bridges. Dev Biol, 305(2), 389–396. doi:10.1016/j.ydbio.2007.02.025

Greenbaum, M. P., Yan, W., Wu, M. H., Lin, Y. N., Agno, J. E., Sharma, M., … Matzuk, M. M. (2006). TEX14 is essential for intercellular bridges and fertility in male mice. Proc Natl Acad Sci U S A, 103(13), 4982–4987. doi:10.1073/pnas.0505123103

Haglund, K., Nezis, I. P., Lemus, D., Grabbe, C., Wesche, J., Liestol, K., … Stenmark, H. (2010). Cindr interacts with anillin to control cytokinesis in Drosophila melanogaster. Curr Biol, 20(10), 944–950. doi:10.1016/j.cub.2010.03.068

Haglund, K., Nezis, I. P., & Stenmark, H. (2011). Structure and functions of stable intercellular bridges formed by incomplete cytokinesis during development. Commun Integr Biol, 4(1), 1–9. doi:10.4161/cib.4.1.13550

Hall, D. H., Winfrey, V. P., Blaeuer, G., Hoffman, L. H., Furuta, T., Rose, K. L., … Greenstein, D. (1999). Ultrastructural features of the adult hermaphrodite gonad of Caenorhabditis elegans: relations between the germ line and soma. Dev Biol, 212(1), 101–123. doi:10.1006/dbio.1999.9356

Hirsh, D., Oppenheim, D., & Klass, M. (1976). Development of the reproductive system of Caenorhabditis elegans. Dev Biol, 49(1), 200–219.

Hu, C. K., Coughlin, M., & Mitchison, T. J. (2012). Midbody assembly and its regulation during cytokinesis. Mol Biol Cell, 23(6), 1024–1034. doi:10.1091/mbc.E11-08-0721

Hubbard, E. J., & Greenstein, D. (2005). Introduction to the germ line. WormBook, 1–4. doi:10.1895/wormbook.1.18.1

Jantsch-Plunger, V., Gonczy, P., Romano, A., Schnabel, H., Hamill, D., Schnabel, R., … Glotzer, M. (2000). CYK-4: A Rho family gtpase activating protein (GAP) required for central spindle formation and cytokinesis. J Cell Biol, 149(7), 1391–1404.

Jenkins, V. K., Timmons, A. K., & McCall, K. (2013). Diversity of cell death pathways: insight from the fly ovary. Trends Cell Biol, 23(11), 567–574. doi:10.1016/j.tcb.2013.07.005

Keil, W., Kutscher, L. M., Shaham, S., & Siggia, E. D. (2017). Long-Term High-Resolution Imaging of Developing C. elegans Larvae with Microfluidics. Dev Cell, 40(2), 202–214. doi:10.1016/j.devcel.2016.11.022

King, R. C., & Mills, R. P. (1962). Oogenesis in adult Drosophila. XI. Studies of some organelles of the nutrient stream in egg chambers of D. melanogaster and D. willistoni. Growth, 26, 235–253.

Kolotuev, I. (2014). Positional correlative anatomy of invertebrate model organisms increases efficiency of TEM data production. Microsc Microanal, 20(5), 1392–1403. doi:10.1017/S1431927614012999

Kolotuev, I., Schwab, Y., & Labouesse, M. (2009). A precise and rapid mapping protocol for correlative light and electron microscopy of small invertebrate organisms. Biol Cell, 102(2), 121–132. doi:10.1042/BC20090096

Lei, L., & Spradling, A. C. (2016). Mouse oocytes differentiate through organelle enrichment from sister cyst germ cells. Science, 352(6281), 95–99. doi:10.1126/science.aad2156

Lekomtsev, S., Su, K. C., Pye, V. E., Blight, K., Sundaramoorthy, S., Takaki, T., … Petronczki, M. (2012). Centralspindlin links the mitotic spindle to the plasma membrane during cytokinesis. Nature, 492(7428), 276–279. doi:10.1038/nature11773

Lewellyn, L., Carvalho, A., Desai, A., Maddox, A. S., & Oegema, K. (2011). The chromosomal passenger complex and centralspindlin independently contribute to contractile ring assembly. J Cell Biol, 193(1), 155–169. doi:10.1083/jcb.201008138

Lints, R., & Hall, D. H. (2009). Reproductive system, germ line. WormAtlas. doi:doi:10.3908/wormatlas.1.23

Loria, A., Longhini, K. M., & Glotzer, M. (2012). The RhoGAP domain of CYK-4 has an essential role in RhoA activation. Curr Biol, 22(3), 213–219. doi:10.1016/j.cub.2011.12.019

Maddox, A. S., Habermann, B., Desai, A., & Oegema, K. (2005). Distinct roles for two C. elegans anillins in the gonad and early embryo. Development, 132(12), 2837–2848. doi:10.1242/dev.01828

Mierzwa, B., & Gerlich, D. W. (2014). Cytokinetic abscission: molecular mechanisms and temporal control. Dev Cell, 31(5), 525–538. doi:10.1016/j.devcel.2014.11.006

Mikeladze-Dvali, T., von Tobel, L., Strnad, P., Knott, G., Leonhardt, H., Schermelleh, L., & Gonczy, P. (2012). Analysis of centriole elimination during C. elegans oogenesis. Development, 139(9), 1670–1679. doi:10.1242/dev.075440

Minestrini, G., Mathe, E., & Glover, D. M. (2002). Domains of the Pavarotti kinesin-like protein that direct its subcellular distribution: effects of mislocalisation on the tubulin and actin cytoskeleton during Drosophila oogenesis. J Cell Sci, 115(Pt 4), 725–736.

Mishima, M., Kaitna, S., & Glotzer, M. (2002). Central spindle assembly and cytokinesis require a kinesin-like protein/RhoGAP complex with microtubule bundling activity. Dev Cell, 2(1), 41–54.

Nishimura, Y., & Yonemura, S. (2006). Centralspindlin regulates ECT2 and RhoA accumulation at the equatorial cortex during cytokinesis. J Cell Sci, 119(Pt 1), 104–114. doi:10.1242/jcs.02737

Odell, G. M., & Foe, V. E. (2008). An agent-based model contrasts opposite effects of dynamic and stable microtubules on cleavage furrow positioning. J Cell Biol, 183(3), 471–483. doi:10.1083/jcb.200807129

Pavicic-Kaltenbrunner, V., Mishima, M., & Glotzer, M. (2007). Cooperative assembly of CYK-4/MgcRacGAP and ZEN-4/MKLP1 to form the centralspindlin complex. Mol Biol Cell, 18(12), 4992–5003. doi:10.1091/mbc.E07-05-0468

Pepling, M. E. (2016). DEVELOPMENT. Nursing the oocyte. Science, 352(6281), 35–36. doi:10.1126/science.aaf4943

Powers, J., Bossinger, O., Rose, D., Strome, S., & Saxton, W. (1998). A nematode kinesin required for cleavage furrow advancement. Curr Biol, 8(20), 1133–1136.

Raich, W. B., Moran, A. N., Rothman, J. H., & Hardin, J. (1998). Cytokinesis and midzone microtubule organization in Caenorhabditis elegans require the kinesin-like protein ZEN-4. Mol Biol Cell, 9(8), 2037–2049.

Robinson, D. N., & Cooley, L. (1997). Genetic analysis of the actin cytoskeleton in the Drosophila ovary. Annu Rev Cell Dev Biol, 13, 147-170.doi:10.1146/annurev.cellbio.13.1.147

Schmutz, C., Stevens, J., & Spang, A. (2007). Functions of the novel RhoGAP proteins RGA-3 and RGA-4 in the germ line and in the early embryo of C. elegans. Development, 134(19), 3495–3505. doi:10.1242/dev.000802

Schonegg, S., Constantinescu, A. T., Hoege, C., & Hyman, A. A. (2007). The Rho GTPase-activating proteins RGA-3 and RGA-4 are required to set the initial size of PAR domains in Caenorhabditis elegans one-cell embryos. Proc Natl Acad Sci U S A, 104(38), 14976–14981. doi:10.1073/pnas.0706941104

Seidel, H. S., Ailion, M., Li, J., van Oudenaarden, A., Rockman, M. V., & Kruglyak, L. (2011). A novel sperm-delivered toxin causes late-stage embryo lethality and transmission ratio distortion in C. elegans. PLoS Biol, 9(7), e1001115. doi:10.1371/journal.pbio.1001115

Severson, A. F., Hamill, D. R., Carter, J. C., Schumacher, J., & Bowerman, B. (2000). The aurora-related kinase AIR-2 recruits ZEN-4/CeMKLP1 to the mitotic spindle at metaphase and is required for cytokinesis. Curr Biol, 10(19), 1162–1171.

Starck, J., & Brun, J. (1977). [Autoradiographic localization of RNA synthesis in vitro during oogenesis in Parascaris equorum]. C R Acad Sci Hebd Seances Acad Sci D, 284(14), 1341–1344.

Sun, L., Guan, R., Lee, I. J., Liu, Y., Chen, M., Wang, J., … Chen, Z. (2015). Mechanistic insights into the anchorage of the contractile ring by anillin and Mid1. Dev Cell, 33(4), 413–426. doi:10.1016/j.devcel.2015.03.003

Vale, R. D., Spudich, J. A., & Griffis, E. R. (2009). Dynamics of myosin, microtubules, and Kinesin-6 at the cortex during cytokinesis in Drosophila S2 cells. J Cell Biol, 186(5), 727–738. doi:10.1083/jcb.200902083

Weber, J. E., & Russell, L. D. (1987). A study of intercellular bridges during spermatogenesis in the rat. Am J Anat, 180(1), 1–24. doi:10.1002/aja.1001800102

White, E. A., & Glotzer, M. (2012). Centralspindlin: at the heart of cytokinesis. Cytoskeleton (Hoboken), 69(11), 882–892. doi:10.1002/cm.21065

Wolfe, B. A., Takaki, T., Petronczki, M., & Glotzer, M. (2009). Polo-like kinase 1 directs assembly of the HsCyk-4 RhoGAP/Ect2 RhoGEF complex to initiate cleavage furrow formation. PLoS Biol, 7(5), e1000110. doi:10.1371/journal.pbio.1000110

Wolke, U., Jezuit, E. A., & Priess, J. R. (2007). Actin-dependent cytoplasmic streaming in C. elegans oogenesis. Development, 134(12), 2227–2236. doi:10.1242/dev.004952

Zanin, E., Desai, A., Poser, I., Toyoda, Y., Andree, C., Moebius, C., Oegema, K. (2013). Aconserved RhoGAP limits M phase contractility and coordinates with microtubule asters to confine RhoA during cytokinesis. Dev Cell, 26(5), 496–510. doi:10.1016/j.devcel.2013.08.005

Zhang, D., & Glotzer, M. (2015). The RhoGAP activity of CYK-4/MgcRacGAP functions non-canonically by promoting RhoA activation during cytokinesis. Elife, 4. doi:10.7554/eLife.08898

Zhou, K., Rolls, M. M., & Hanna-Rose, W. (2013). A postmitotic function and distinct localization mechanism for centralspindlin at a stable intercellular bridge. Dev Biol, 376(1), 13–22. doi:10.1016/j.ydbio.2013.01.020

